# Regulation of FGF/MAPK signaling at the primary cilium base controls the proliferation-neurogenesis balance in human cerebellar organoids

**DOI:** 10.1101/2025.04.14.648790

**Authors:** Ludovica Brunetti, Antonia Wiegering, Isabelle Anselme, Minh Ngoc Vu, Lidia Pollara, Marie Grappin, Martin Catala, Olivier Saulnier, Christophe Antoniewski, Enza Maria Valente, Sylvie Schneider-Maunoury, Christine Vesque

## Abstract

The primary cilium acts as a specialized signaling compartment that coordinates multiple developmental pathways, yet its role in regulating signaling activity during human brain development remains poorly understood. Biallelic mutations in ciliary genes cause neurodevelopmental ciliopathies such as Joubert syndrome, which are characterized by cerebellar hypoplasia and dysplasia. Here, we generated human cerebellar organoids carrying null or patient-derived mutations in RPGRIP1L, a ciliary transition zone gene associated with Joubert syndrome, to investigate how ciliary dysfunction alters human cerebellar neurogenesis. While control organoids robustly express markers of cerebellar glutamatergic and GABAergic lineages including Purkinje cells, RPGRIP1L-deficient organoids display a consistent and severe reduction in Purkinje cell markers, accompanied by impaired neurogenesis and increased progenitor proliferation. These defects coincide with prolonged overactivation of the FGF/MAPK signaling pathway. We further show that the MAPK effector pMEK1/2 localizes at the base of primary cilia, where its levels are significantly increased in RPGRIP1L-deficient cerebellar progenitors. Pharmacological inhibition of FGF receptors reduces pMEK1/2 activation at cilia base and rescues both the proliferation-neurogenesis imbalance and Purkinje lineage defects, without restoring the underlying ciliary abnormalities. Together, our findings identify the primary cilium as a compartment that refines FGF/MAPK signaling in human cerebellar progenitors, and support a model in which RPGRIP1L restrains FGF pathway activity to promote neurogenesis. These results provide a mechanism linking ciliary dysfunction to altered cerebellar development and provide new insight into the developmental origin of cerebellar impairment in neurodevelopmental ciliopathies

## INTRODUCTION

Primary cilia are microtubular sensory organelles that act as signaling hubs coordinating multiple developmental pathways, including HH, WNT, PDGF and FGF signaling [1–3]. To ensure proper signaling, their composition is tightly controlled by the transition zone, a molecular gate formed by protein complexes including the MKS, NPHP (NPHP1, NPHP4, NPHP8/RPGRIP1L), and CEP290 modules [4,5]. Disruption of ciliary structure or function causes a broad spectrum of developmental disorders collectively known as ciliopathies [1]. Among them, Joubert syndrome (JBTS) is a rare neurodevelopmental disease caused by biallelic mutations in one of over 40 associated genes, all encoding proteins involved in primary cilia structure or function [6,7]. Clinically, JBTS patients present with ataxia, ocular motor defects, developmental delay, and variable levels of cognitive impairment [8]. Despite the high genetic and phenotypic variability, all JBTS patients share the diagnostic “molar tooth sign” on axial brain imaging, resulting from specific cerebellar and brainstem malformations, most notably the hypoplasia/dysplasia of the vermis – the medial cerebellar structure [6,9].

Cerebellar dysfunction is a common feature of several developmental disorders, associated with both motor disease and cognitive impairment [10]. The cerebellum originates from dorsal hindbrain rhombomere 1 (r1) under inductive cues from the isthmic organizer at the midbrain–hindbrain boundary, where fibroblast growth factor 8 (FGF8) expression defines the early cerebellar territory [11–13]. Two distinct germinal zones subsequently form: the ventricular zone (VZ), which produces inhibitory GABAergic neurons such as Purkinje, Golgi, and basket cells [14], and the rhombic lip (RL), which generates excitatory glutamatergic lineages including granule neurons (GNs) and unipolar brush cells [15,16]. In humans, Purkinje cell neurogenesis occurs between post-conception weeks (PCW) 8–10 [17,18], whereas granule cell progenitors (GCPs) proliferate extensively around 26 PCW [19]. Coordinated proliferation and differentiation of these populations are critical for establishing cerebellar size, morphology, and function [20].

In the cerebellum, ciliogenesis is temporally regulated to ensure appropriate responsiveness to SHH and other signals [21–23]. In mice, a loss of cilia in the brain causes cerebellar hypoplasia and foliation defects due to loss of SHH-dependent GCP proliferation [24,25], while mouse mutants for JBTS genes — including Rpgrip1l [26], Ahi1 [27], Tmem67 [28], Jbts17 (CPLANE1) [29], Talpid3 [30], Arl13b [31] and Cep290 [27] — exhibit variable cerebellar phenotypes and divergent effects on HH and WNT signaling. In humans, the analysis of JBTS fetal samples with a confirmed diagnosis of cerebellar vermis hypoplasia revealed a reduction of GCP proliferation associated with a lower expression of HH targets, with no differences between hemispheres and vermis. These findings suggest that the specific vermis hypoplasia observed in JBTS patients precedes and is independent of GCP proliferation defects, leaving its primary origin unresolved [32]. Overall, the pathogenetic mechanisms linking cilia dysfunction to JBTS neurodevelopmental defects remain largely unknown and may be of diverse origins.

This leaves us with two main open questions: i) what are the early cellular and molecular processes affected by ciliary gene depletion during human cerebellar development? ii) can some of these alterations account for the neurodevelopmental defects underlying cerebellar vermis hypo/dysplasia in JBTS patients? In order to address these questions, we turned to human iPSC-derived cerebellar organoids, recently established as promising approaches to study human cerebellar development and neurodevelopmental diseases [33–40]. By recapitulating developmental pathways, these approaches are particularly well suited to study early steps of cerebellar development. Moreover, emerging evidence indicates that primary cilia themselves display human-specific features and can contribute to species- specific aspects of neural progenitor behavior and brain development [41,42]. In a previous work we demonstrated significant differences concerning cilia formation and function in human and mouse spinal organoids in the absence of JBTS proteins (RPGRIP1L and TMEM67) [42], further emphasizing the importance of using human-based systems to investigate the developmental origin of cilia-related cerebellar defects.

Here, we employ human iPSC-derived cerebellar organoids to investigate the role of cilia and of the ciliary protein RPGRIP1L in cerebellar development. RPGRIP1L encodes a scaffolding protein essential for the integrity of the ciliary transition zone [43,44] and HH pathway activity [45,46]. Mutations in RPGRIP1L can cause both JBTS and the more severe Meckel syndrome in humans [26], and are responsible for severe neurodevelopmental defects in mouse models [47]. Using organoids derived from JBTS patient iPSCs carrying biallelic RPGRIP1L mutations and CRISPR-engineered knockouts, we identify the primary cilium base as a compartment for FGF/MAPK signaling regulation in human cerebellar progenitors. RPGRIP1L deficiency results in early ciliary abnormalities, aberrant FGF/MAPK signaling activation with increased accumulation of phosphorylated MEK1/2 at the ciliary base, excessive progenitor proliferation and impaired neurogenesis. These defects are rescued by pharmacological inhibition of the FGF pathway after midbrain-hindbrain boundary induction.

Our findings reveal a causal link between RPGRIP1L dysfunction, dysregulated FGF signaling, and disrupted cerebellar neurogenesis, providing mechanistic insight into the role of primary cilia in cerebellar development and the origin of cerebellar malformations in JBTS.

## RESULTS

### Cerebellar organoids generated from human induced pluripotent stem cells recapitulate early steps of cerebellar development

To unravel molecular mechanisms and biological processes that may be responsible for cerebellar defects in JBTS pathogenesis, we used organoids to model early stages of human cerebellar development in vitro. We optimized published protocols for generating cerebellar organoids from human induced pluripotent stem cells (hiPSCs) [33,35]. To induce neural commitment, mid-hindbrain boundary establishment and tissue self-organization, embryoid bodies were sequentially treated with fibroblast growth factor 2 (FGF2), insulin, TGFß/SMAD inhibitor (SB431542), FGF19 and stromal cell-derived factor 1 (SDF1) (Fig. 1A). Since dynamic culture conditions have been shown to favor faster cerebellar commitment [48], embryoid bodies were cultured under orbital agitation from day 7 onwards.

**FIGURE 1.**
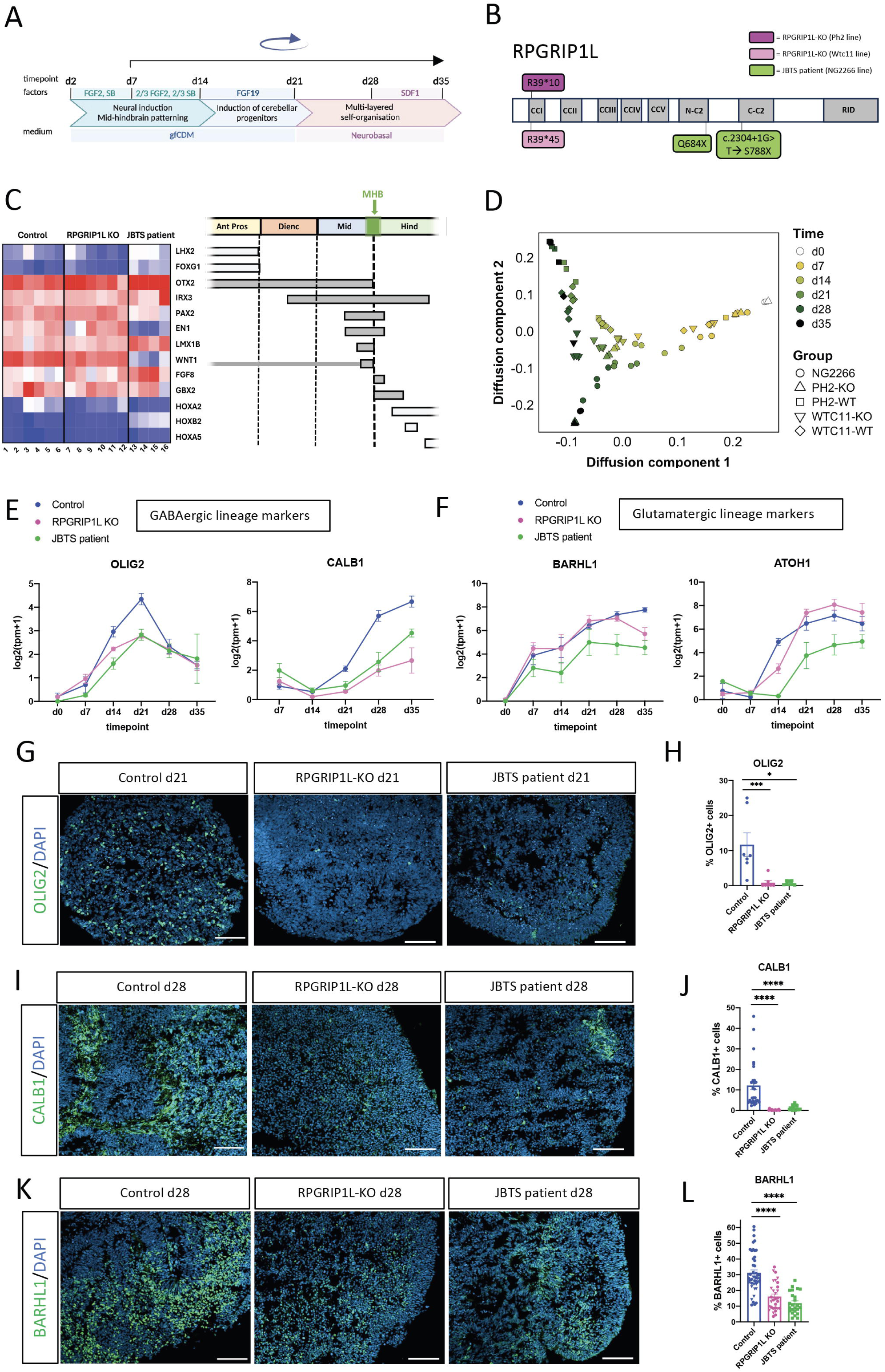
RPGRIP1L-deficient cerebellar organoids present a severe reduction of Purkinje cell markers and a mild decrease of glutamatergic lineage markers (A) Diagram of the 3D cerebellar differentiation protocol. Time-line and drugs (SB: SMAD antagonists SB-431542; FGF2: Fibroblast growth factor 2; FGF19: Fibroblast growth factor 19; SDF1: stromal cell-derived factor 1) are indicated on top, culture media at the bottom. The blue circular arrow on top indicates orbital rotation. Protocol modified after [35] (B) Schematic representation of the RPGRIP1L protein and relative position of the mutations present in hiPSC lines. In both RPGRIP1L KO lines (Ph2 in purple, Wtc11 in pink) homozygous frameshift mutations result in premature stop codon before the first coiled- coil (CC) domain. JBTS patient (NG2266 in green) variants p.Q684X and c.2304+1G>T result in stop codons between the first and the second C2 domain. (C) Brain marker gene expression in cerebellar organoids at day 7. Heatmap based on the normalized expression values of selected brain marker genes. Z score was calculated on the log2(tpm+1) data from the bulk RNASeq analysis. Right: scheme of the gene expression domains in the E9.5 mouse brain. MHB: midbrain-hindbrain boundary. Thinner box for Wnt1: expression restricted to the roof plate. See supplementary Table 1 for number of independent experimental replicates (N) and sample numbering. (D) Diffusion map of the RNAseq data of all cell lines at different days of differentiation. (E, F) Graphs showing temporal gene expression analysis of GABAergic lineage markers (OLIG2 and CALB1) (E) and glutamatergic lineage markers (BARHL1 and ATOH1) (F) in control (blue), RPGRIP1L KO (magenta), and JBTS patient-derived (green) organoids throughout differentiation. Normalized gene expression (Log2(tpm+1)) values from bulk RNASeq analysis are displayed as mean ± SEM. More than 20 organoids were analyzed per condition per experiment. For control and RPGRIP1L KO, data were obtained from both Ph2 and Wtc11 lines. Day 0 (hiPSCs): N=2 for controls, N=2 for RPGRIP1L KO, N=1 for JBTS patient line; day 7-35 organoids: see Supplementary Table 2 for number of independent experimental replicates. (G) Immunofluorescence for OLIG2 (Purkinje lineage progenitors) on cryosections from control, RPGRIP1L KO and JBTS patient organoids at day 21. (H) Graph showing the % of OLIG2+ over DAPI+ nuclei. Each dot represents an organoid section. Several sections from different organoids were analyzed per condition (N=1). Data are shown as mean ± SEM. Asterisks denote statistical significance according to One-Way ANOVA Kruskal-Wallis test with Dunn’s correction (***P<0.0002 and ****P< 0.0001). (I) Immunofluorescence for CALB1 on cryosections from day 28 control, RPGRIP1L KO and JBTS patient organoids. (J) Quantification of the % of CALB1+ cells over DAPI+ nuclei, on > 15 sections from different organoids per experiment per condition (control and RPGRIP1L KO: Ph2 line N=2 – round dots; JBTS patient: NG2266 line N=2 – squared dots). Data are shown as mean ± SEM. Asterisks denote statistical significance according to One-Way ANOVA Kruskal-Wallis test with Dunn’s correction (****P<0.0001). (K) Immunofluorescence for BARHL1 on cryosections from day 28 control, RPGRIP1L KO and JBTS patient organoids. (L) Quantification of the % of BARHL1+ over DAPI+ nuclei, in > 15 sections from different organoids per experiment per condition (control and RPGRIP1L KO N=3: Ph2 line N=2 – round dots, Wtc11 line N=1 – triangular dots; JBTS patient: NG2266 line N=2 – squared dots). Data are shown as mean ± SEM. Asterisks denote statistical significance according to One-Way ANOVA Kruskal-Wallis test with Dunn’s correction (****P<0.0001). N: number of independent experimental replicates. Scale bars: 100 µm in F, H and J.

Immunostaining analysis on control organoid cryosections showed clusters of neural progenitors (SOX2^+^) and early-born neurons (HUC/D^+^), already present at day 14 (Supplementary Fig. 1A), progressively expanding and self-organizing (Supplementary Fig. 1A-C). By day 28, early-born neurons are primarily located in deep regions of the organoid tissue, adjacent to an outer layer of neural progenitors organized either at the organoid periphery or into small internal rosettes also positive for the apical junction marker ZO1 (Supplementary Fig. 1C, E). At this stage, the cerebellar VZ marker KIRREL2 was present in SOX2-positive rosettes and enriched at the apical surface (Supplementary Fig. 1D), while the glutamatergic progenitor marker BARHL1 and the GABAergic neuron marker CALB1 segregated in separated domains within the organoid tissue, with few cells intermingled (Supplementary Fig. 1F).

To assess the time-course of cerebellar cell type production in our experimental settings, we performed bulk RNAseq analysis on cerebellar organoids derived from two independent control hiPSC lines with different genetic backgrounds, PCIi033-A (Ph2) and UCSFi001-A (Wtc11) at five timepoints of the differentiation protocol (day 7, 14, 21, 28, 35) (see Supplementary Excel File 1 for normalized gene expression data). First, we validated that the control organoids faithfully recapitulated early stages of cerebellar development, with the presence of midbrain, hindbrain and mid-hindbrain boundary markers and the absence of anterior forebrain and posterior hindbrain markers (Figure 1C). Concerning cerebellar differentiation, the order of appearance of GABAergic Purkinje lineage markers reflected the in vivo maturation stages of this cell population, with the progenitor marker OLIG2 and the early post-mitotic marker SKOR2 peaking at day 21, followed by the more mature Purkinje marker CALB1, strongly expressed between day 28 and 35 (Supplementary Fig 1G). As for the glutamatergic lineage, the RNA levels of progenitor markers were gradually increased over time, first appearing at day 7 for PAX6 and BARHL1, and at day 14 for ATOH1 (Supplementary Fig. 1H). Of note, the mature GN marker, GABRA6, was not detected over the 35 days short-term organoid culture, consistent with its post-natal onset in humans [18,49] (Supplementary Fig. 1I). Given the well-established role of the SHH pathway in GCP pool expansion during cerebellar development [50,25], we also examined SHH ligand expression. However, almost no SHH mRNA was detected over the 35 days of cerebellar differentiation (Supplementary Fig. 1J).

Altogether these data provide strong evidence for self-organization of a cerebellar primordium and successful specification of the two main cerebellar lineages following our differentiation protocol. They also show that this 35-days differentiation protocol models early cerebellar developmental stages, preceding SHH endogenous expression by Purkinje cells.

### RPGRIP1L-deficient cerebellar organoids present a severe reduction of PC markers and a milder decrease in the glutamatergic lineage

To compare the differentiation process between healthy and JBTS pathological conditions, we used two independent RPGRIP1L KO hiPSC lines – generated in the two different genetic backgrounds previously used for control organoid analysis (Ph2 and Wtc11) [42] – both producing early truncated proteins within the first coiled coil domain (Fig. 1B). In parallel we also analyzed a hiPSC line derived from a severely affected JBTS patient (NG2266) carrying two pathogenic variants in the RPGRIP1L gene (p.Q684X/c.2304+1G>T) [51] (Fig. 1B). Both variants of this patient are predicted by VarSome to be null variants and are classified as pathogenic following the pathogenicity criterium “very strong” (PVS1) accordingly to the ACMG (American College of Medical Genetics and Genomics) standards and guidelines [52] and encode truncated proteins missing the second C2 domain and C-terminal RPGR- Interacting domain (RID) of the protein (see Methods and Supplementary Fig. 8). We generated cerebellar organoids from both RPGRIP1L KO hiPSC lines (Ph2 and Wtc11) in parallel with their isogenic controls, and from the JBTS patient-derived hiPSC line (NG2266).

Using a specific antibody raised against the human RPGRIP1L RID domain, we showed the presence of RPGRIP1L protein at its subcellular localization at the ciliary transition zone in control organoids (Supplementary Fig. 2A, A’). On the contrary, no signal was detected in cilia from both RPGRIP1L KO and JBTS patient-derived organoids (Supplementary Fig. 2B-C’).

Bulk RNAseq analysis was performed on hiPSC cells and on cerebellar organoids collected every seven days from day 7 to day 35 from several independent experiments (see Supplementary Excel File 1 for normalized gene expression data). We showed that both RPGRIP1L KO and JBTS patient-derived organoids at day 7 expressed mid-hindbrain markers, while being negative for anterior forebrain and caudal hindbrain markers (Fig 1C), thus showing proper specification of their early anteroposterior identity. However, JBTS patient- derived organoids displayed lower levels of WNT1 and EN1 than controls. A diffusion map of the RNAseq data illustrated i) the distribution of samples along the DC1 axis according to increasing culture time, reflecting their progressive differentiation, and ii) after day 14, the segregation along the DC2 axis of the control samples (Ph2 and wtc11) on one side and RPGRIP1L-deficient (both RPGRIP1L-KO and JBTS patient-derived) organoid samples on the other side, indicating an increasing divergence in their molecular identities (Fig. 1D). From day 14 onwards, transcriptome analysis revealed a consistent and severe reduction of numerous Purkinje cell markers (OLIG2, SKOR2, CALB1, FOXP2, PDE1C, ITPR1) in both RPGRIP1L KO and JBTS patient-derived organoids compared to controls (Fig. 1E, Supplementary Fig. 2D). Quantitative PCR analysis corroborated these results showing a significant decrease of OLIG2, SKOR2 and CALB1 markers in RPGRIP1L KO organoids derived from both hiPSC lines and in JBTS patient-derived organoids, compared to controls (Supplementary Fig. 2E).

In contrast, the expression levels of the glutamatergic lineage markers ATOH1 and BARHL1 were not consistently affected across RPGRIP1L-deficient conditions compared to controls (Figure 1F). This was confirmed by quantitative PCR analysis where the only significant defects were a decrease in BARHL1 expression in JBTS patient-derived organoids and an increase in ATOH1 expression in RPGRIP1L KO organoids at d35 (Supplementary Fig. 2F).

In order to strengthen these findings, we assayed protein expression of both Purkinje lineage markers (OLIG2, CALB1) and glutamatergic lineage markers (ATOH1, BARHL1) by quantifying the number of positive cells after immunostaining on organoid cryosections. Each marker was analyzed at the timepoint of its highest expression in control organoids, as determined by bulk RNA-seq data: day 21 for OLIG2 and day 28 for CALB1, ATOH1, and BARHL1. OLIG2- and CALB1-positive cells were barely detected in RPGRIP1L KO and JBTS patient-derived organoids, while strong and widespread labeling was observed in control organoids (Fig. 1G- J, Supplementary Fig. 2I). Regarding the glutamatergic lineage, the number of BARHL1 positive cells was reduced by 50% in RPGRIP1L KO and JBTS patient-derived organoids, along with a decrease in the average labeling intensity of individual cells (Fig. 1K,L, Supplementary Fig. 2J). The number of ATOH1 positive cells was only reduced in JBTS patient organoids (Supplementary Fig. 2G, H) whereas the intensity of ATOH1 labelling was reduced by ∼50% both in JBTS patient and RPGRIP1L KO conditions (Supplementary Fig. 2K).

Overall, these data indicate a severe and significant impairment in the specification and production of the Purkinje lineage in all RPGRIP1L-deficient conditions from day 21 onwards, whereas glutamatergic lineage specification appeared to be modestly affected, mainly at protein level from day 28.

### Increased proportion of early neural progenitors in RPGRIP1L KO and JBTS patient cerebellar organoids

We assessed whether these molecular alterations observed upon RPGRIP1L impairment might be preceded by earlier defects in tissue organization. The first obvious morphological alteration under brightfield observation was a size increase of RPGRIP1L KO and JBTS patient organoids compared to controls, from day 21 onward (Fig. 2A). Organoid area was measured every 7 days along the differentiation protocol in several independent experiments. At day 28 and 35, both RPGRIP1L KO and JBTS patient-derived organoids presented a significant area increase compared to controls (3.3-fold for RPGRIP1L KO and 2.3-fold for JBTS patient- derived organoids over controls at day 35) (Fig. 2B).

**FIGURE 2.**
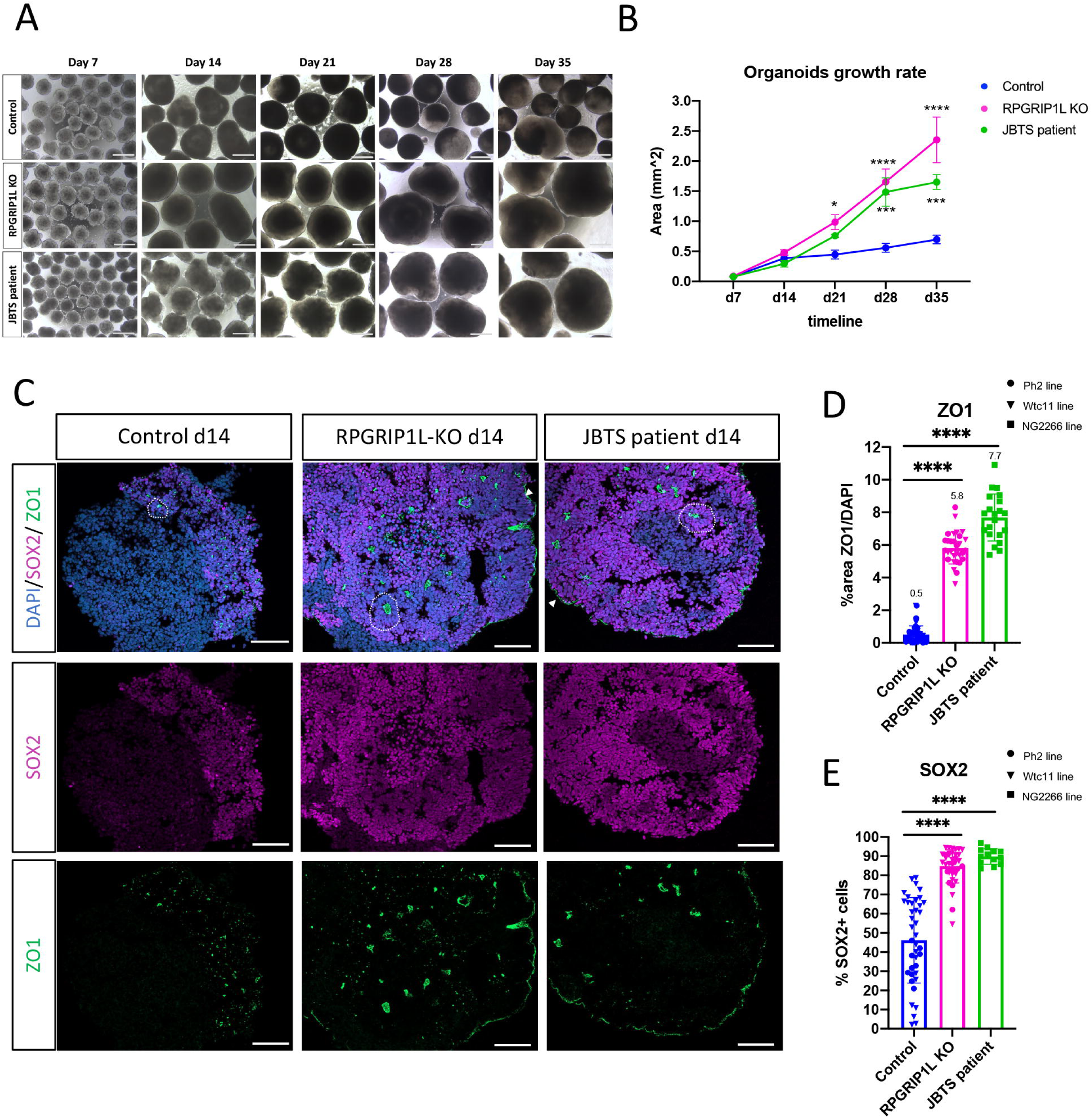
Organoid size and area of SOX2 /ZO1 progenitors are increased in RPGRIP1L- deficient cerebellar organoids. (A) Representative bright field images of cerebellar organoids at the indicated time points. (B) Graph representing the organoid growth rate, measured every 7 days from day 7 to day 35. Data are shown as the mean ± SEM of average organoid area from several independent experiments (N=3 Ph2 line, N=3 Wtc11 line, N=3 NG2266 line, n=10-60 organoids analyzed per experiment per condition at each timepoint). For control and RPGRIP1L KO conditions, data from Ph2 and Wtc11 lines were pooled. Asterisks denote statistical significance according to Two-Way ANOVA test (*P<0.033, ***P<0.0002, ****P<0.0001). (C) Immunofluorescence for SOX2 neural progenitor marker (magenta) and ZO1 apical marker (green) on cryosections from control, RPGRIP1L KO and JBTS patient organoids at day 14. Examples of small neural rosettes are circled by dotted lines. (D) Quantification of the ZO1+ area over the DAPI+ area in sections from control, RPGRIP1L KO and JBTS patient organoids at day 14. Each dot represents an organoid section. Several sections from different organoids were analyzed per condition: control and RPGRIP1L KO: N=2 (Ph2 N=1 – round dots, Wtc11 N=1 – triangular dots), JBTS patient: NG2266 N=1 – squared dots. Mean ± SEM are indicated. Asterisks: statistical significance according to One-Way ANOVA Kruskal-Wallis test with Dunn’s correction (****P< 0.0001) (E) Graph showing the % of SOX2^+^ cells in sections from control, RPGRIP1L KO and JBTS patient organoids at day 14. Each dot represents an organoid section. Several sections from different organoids were analyzed per condition: control and RPGRIP1L KO: N=3 (Ph2 N=1 – round dots, Wtc11 N=2 – triangular dots), JBTS patient: NG2266 N=1 – squared dots. Mean ± SEM are indicated. Asterisks denote statistical significance according to One-Way ANOVA Kruskal-Wallis test with Dunn’s correction (****P< 0.0001). N: number of independent experimental replicates. Scale bars: 500 µm in A and 100 µm in C.

Moreover, the general tissue organization was already perturbed at day 14, before overt size increase. In control organoids, early polarized neural progenitors (SOX2^+^/ZO1^+^) formed a few small internal rosettes (Fig. 2C left panels). By contrast, RPGRIP1L KO (Ph2 and Wtc11 lines) and JBTS patient-derived organoids exhibited a higher number of internal rosettes. Additionally, in all RPGRIP1L-deficient conditions, polarized neural progenitors (SOX2^+^/ZO1^+^) extended across the entire organoid surface, with their apical side oriented outward, forming a continuous neuroepithelium in contact with the medium (Fig. 2C, middle and right panels).

The area occupied by ZO1 apical surface within the organoid was expanded by 12-fold in RPGRIP1L-KO (Ph2 and Wtc11 lines) and by 15-fold in JBTS patient-derived organoids compared to control condition at day 14 (Fig. 2D). Similarly, the number of SOX2+ neural progenitors was doubled in all RPGRIP1L-deficient conditions compared to controls (Fig. 2E).

### Differential dependence on RPGRIP1L protein among neural progenitors for ciliogenesis

Given the well-established role of RPGRIP1L in building functional cilia [43,53,44] and the fact that apical progenitors of the neural tube are ciliated [54–57], we investigated whether ciliogenesis was affected in human cerebellar organoids. Immunostaining for the axonemal marker INPP5E and for the basal body marker gamma-TUBULIN revealed numerous cilia pointing towards the internal lumen of neural rosettes in both controls (Fig. 3A, A’) and all RPGRIP1L-deficient conditions (Fig. 3B-C’). Some cilia in RPGRIP1L KO and JBTS rosettes displayed a bulbous tip (Fig. 3B’, C’), a condition which was never observed in control rosettes. However, ciliary length was not significantly changed in RPGRIP1L KO rosettes and only mildly decreased in JBTS condition (Fig. 3D).

**FIGURE 3.**
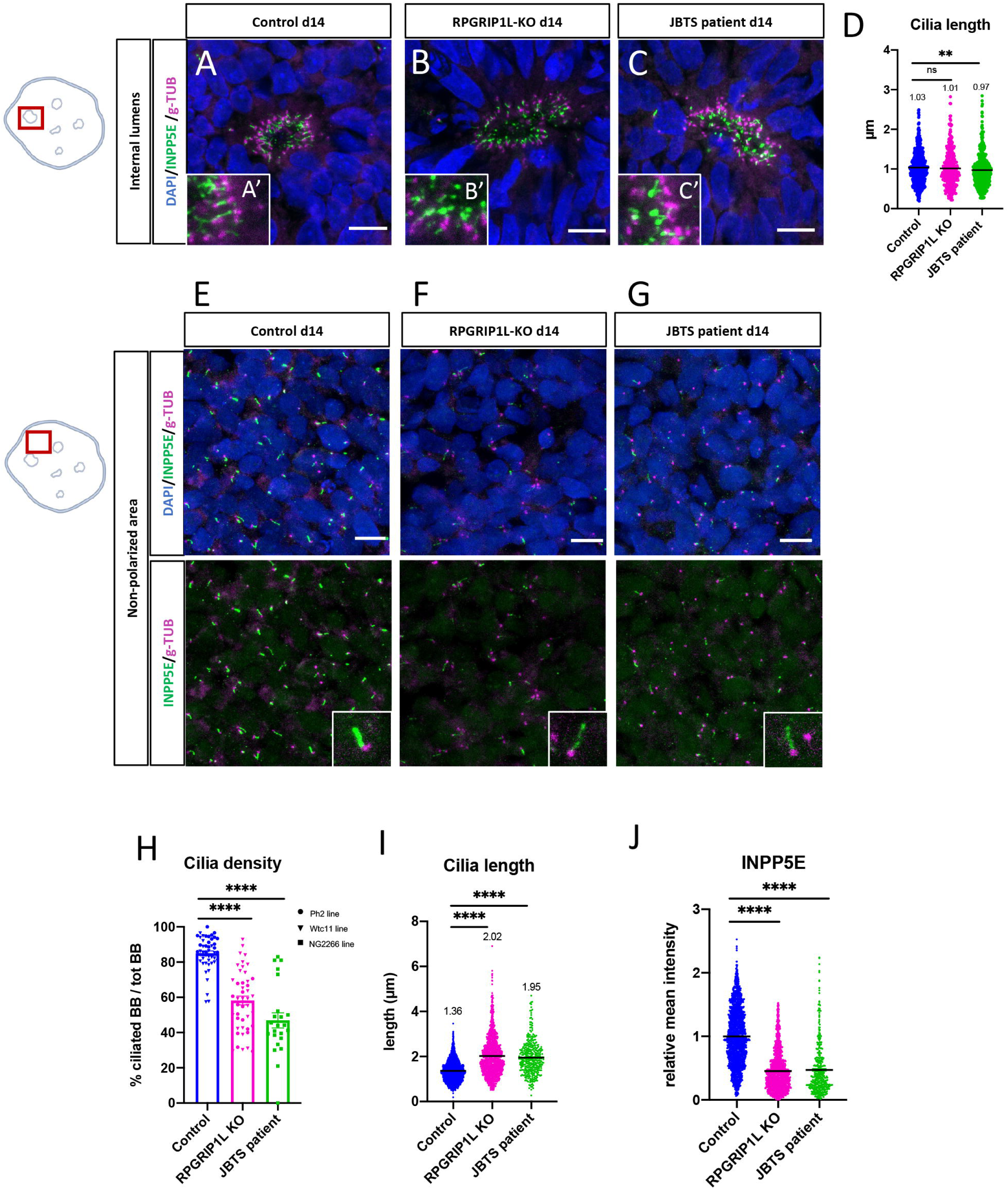
Differential requirement of RPGRIP1L protein for ciliogenesis in apical and non- polarized neural progenitors. (A-C) Immunofluorescence for INPP5E (green) and gamma-TUBULIN (magenta) on sections from control (A), RPGRIP1L KO (B) and JBTS patient (C) organoids at day 14. Scale bar, 10 µm. The schematic on the left represents the position of the scanned regions: i.e. internal lumen of neural rosettes. (D) Graph showing the quantification of ciliary length. Measures for individual cilia are represented and the mean value is indicated. Asterisks: statistical significance according to One-Way ANOVA Kruskal-Wallis with Dunn’s correction (**P < 0.0021). (E-G) Immunofluorescence for INPP5E (green) and gamma-TUBULIN (magenta) on sections from control (E), RPGRIP1L KO (F) and JBTS patient (G) organoids at day 14. Scale bar, 10 µm. The schematic on the left represents the position of the scanned regions: i.e. non-polarized tissue. (H-J) Graphs showing quantifications of ciliary properties in non- polarized areas of control, RPGRIP1L KO and JBTS patient cerebellar organoids. Asterisks denote statistical significance according to One-Way ANOVA Kruskal-Wallis test with Dunn’s correction (**P < 0.0021, ****P < 0.0001). Control and RPGRIP1L KO: N=4 (Wtc11 N=3 – triangular dots, Ph2 N=1 – round dots); JBTS patient: NG2266 N=2 – squared dots. N: number of independent experimental replicates. In (H) Ciliary density is calculated as the % of gamma-TUBULIN+ basal bodies presenting an INPP5E+ axoneme over the total number of basal bodies per field. Each dot on the graph represents one quantified field representative for one organoid section. In (I-J) each dot represents one individual cilium. Average cilia length (I) and average axonemal INPP5E intensity after background subtraction (J) are calculated from >400 cilia per condition measured in >20 organoids from independent experiments and hiPSC lines. Scale bars: 10 µm in A-C and E-G.

In contrast, the analysis of non-polarized regions, characterized by random nuclear orientation and distribution of gamma-TUBULIN+ basal bodies (Fig. 3E-G), revealed a significant reduction in the number of ciliated basal bodies in RPGRIP1L-deficient conditions (58% in RPGRIP1L-KO and 47% in JBTS patient organoids, versus 85% in controls) (Fig. 3H). Moreover, ciliary length increased (Fig. 3I) and INPP5E labeling intensity inside cilia was reduced (Fig. 3J), suggesting defective ciliary gating.

The analysis of two TZ proteins belonging to different multiprotein complexes, NPHP1 and TMEM67, revealed a near-total loss of NPHP1, a member of the NPHP complex, in all cilia of RPGRIP1L KO organoids (Supplementary Fig. 3A), while the MKS complex member TMEM67 was preserved and even slightly more abundant (Supplementary Fig. 3B). Together, our findings suggest that, in cerebellar organoids, RPGRIP1L is necessary for correct TZ formation, as well as for proper ciliogenesis and ciliary content in non-polarized neural progenitors, while it appears dispensable for ciliogenesis in ZO1^+^ early human cerebellar progenitors at day 14. Nevertheless, the observation of numerous bulbous tips on cilia of apical progenitors suggests that they might present an abnormal content, likely to modify their signaling properties.

### RPGRIP1L loss leads to neurogenesis impairment and transcriptional upregulation of FGF pathway genes

In order to identify specific molecular pathways disrupted upon RPGRIP1L depletion we performed a DESeq2 analysis on our bulk RNAseq dataset, comparing both RPGRIP1L KO and JBTS patient-derived organoids with controls at selected timepoints. We aimed at identifying robust and shared phenotypes related to RPGRIP1L deficiency across all genetic backgrounds.

At day 14, many genes appeared deregulated in both RPGRIP1L KO (2146 genes, P adj <0.05, |log2FC|>1) and JBTS patient-derived organoids (4304 genes, P value <0.05, |log2FC|>1) compared to controls (Supplementary Excel Files 2 and 3, respectively). The Volcano plots representing significant DEGs (P value <0.05, |log2FC|>1) for RPGRIP1L KO compared to control (Fig. 4A) or JBTS patient compared to control (Fig. 4B) revealed many genes commonly de-regulated in both RPGRIP1L-deficient conditions. Among the most downregulated genes were neurogenic master genes – NEUROG1, NEUROD1 and NEUROD4 – as well as other genes involved in neurogenesis and early-born neuron functions like HES6 [58,59], GDNF and RELN (Fig. 4A, B). On the contrary, several SOX and HES transcription factors, important to maintain a neural progenitor state, were significantly up-regulated in both RPGRIP1L KO and JBTS patient-derived organoids compared to controls (Fig. 4A, B).

**FIGURE 4.**
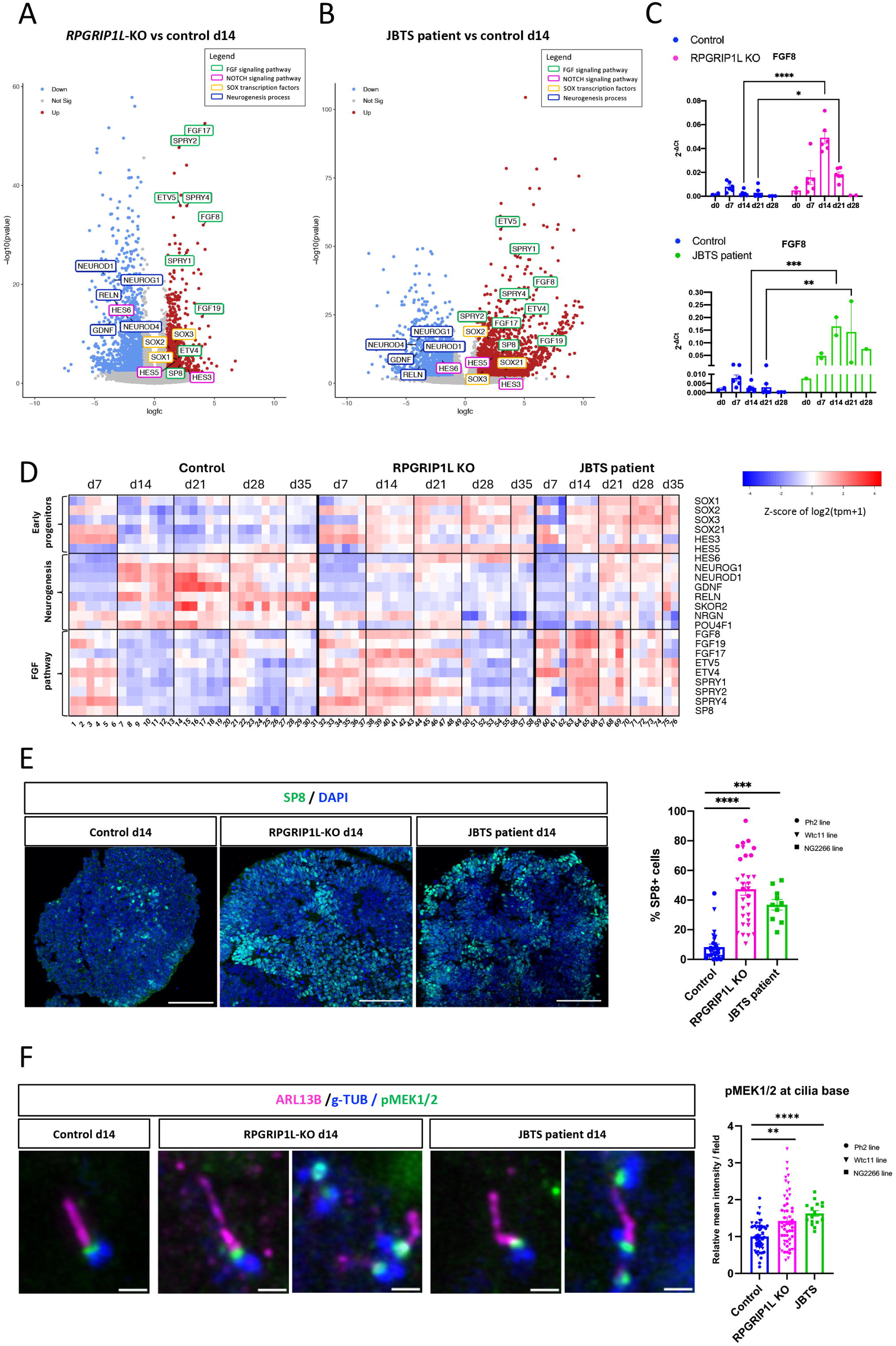
Transcriptome analysis reveals early FGF signaling upregulation and neurogenesis impairment in RPGRIP1L-deficient cerebellar organoids (A, B) Volcano plots representing differentially expressed genes (DEGs) resulting from DESeq2 analysis (only genes with P<0.05; |log_2_FC|>1 are colored) in RPGRIP1L KO versus control condition (A) or in JBTS patient versus control condition (B) at day 14. Selected genes from representative pathways and cellular processes are circled with different colors as indicated in the legend on the top right of the panels. (C) qPCR analysis of FGF8 expression levels on cerebellar organoids at different timepoints; control and RPGRIP1L KO: N=6 (Ph2 N=2, Wtc11 N=4); JBTS patient: N=2. Data are shown as individual values; mean ± SEM are indicated. Asterisks denote statistical significance according to ordinary Two-Way ANOVA performed on control and RPGRIP1L KO samples (*P<0.033, **P<0.0021, ***P<0.0002, ****P < 0.0001). N: number of independent experimental replicates. (D) Heatmap representing normalized RNA expression values of selected DEGs in control and RPGRIP1L- deficient cerebellar organoids over time. >20 organoids per replicate per condition were collected at each timepoint; See supplementary Table 2 for number of independent experiments and sample numbering. (E) Confocal images of immunofluorescence for SP8 (green) on cryosections from control and RPGRIP1L-deficient organoids at day 14. Nuclei are stained with DAPI (blue). Right: graph showing the % of SP8+ over DAPI+ nuclei, counted in > 10 sections from different organoids per condition per experiment. Controls and RPGRIP1L KO: N=3 (Wtc11 N=2 – triangular dots, Ph2 N=1 – circular dots), JBTS: NG2266 N=1 squared dots. Each dot in the graph represents one organoid; mean ± SEM are shown. Asterisks denote statistical significance according to One-Way ANOVA Kruskal-Wallis test with Dunn’s correction (***P<0.0002; ****P<0.0001). Scale bar, 100 µm. N: number of independent experimental replicates. (F) High-resolution AIRYSCAN immunofluorescence images for pMEK1/2 (green), ARL13B (magenta) and gamma-TUBULIN (blue) on cryosections from control and RPGRIP1L-deficient organoids at day 14. Scale bar, 1 µm. Right: graph showing the mean % of pMEK1/2+ over gamma-TUBULIN+ basal bodies, counted in > 10 sections from different organoids per condition per experiment. Controls and RPGRIP1L KO: N=4 (Wtc11 N=2 – triangular dots, Ph2 N=2 – circular dots), JBTS: NG2266 N=2 squared dots. Each dot in the graph represents one section analyzed; mean ± SEM are shown. Asterisks denote statistical significance according to One-Way ANOVA Kruskal-Wallis test with Dunn’s correction (**P<0.0021, ****P<0.0001). N: number of independent experimental replicates.

Most importantly, FGF signaling emerged as a severely disrupted pathway, with many FGF ligands (FGF8, FGF19, FGF17) and downstream targets (ETV4, ETV5, SPRY1-2-4, SP8) strongly up-regulated in both RPGRIP1L KO and JBTS patient-derived organoids (Fig.4A, B). In particular, FGF8 emerged as the top up-regulated FGF ligand gene, with a peak of maximal over-expression at day 14 in RPGRIP1L KO organoids (log2FC=4.07, p adj=7.10E-30) and JBTS patient organoids (log2FC=6.01, p adj=3.24E-32). Real-time quantitative PCR analysis of FGF8 expression confirmed this up-regulation between day 14 and day 21 in RPGRIP1L KO and JBTS patient conditions versus controls and a more exacerbate phenotype in JBTS patient organoids (Fig. 4C). To illustrate the reproducibility of the differentiation protocol as well as the similarity of transcriptional changes between RPGRIP1L KO and JBTS patient-derived organoids, we generated a heatmap representation of normalized RNA expression values for selected genes over time (Fig. 4D). This showed that, in control samples, neurogenesis genes started to be expressed around day 14, concomitant with a downregulation of early progenitor marker genes. In RPGRIP1L-deficient conditions, these changes were delayed or absent. Moreover, while in control organoids FGF signaling peaked at day 7 and was turned off by day 14, in both RPGRIP1L KO and JBTS patient-derived organoids this signaling was upregulated and longer maintained (until day 21). Overall, these data suggest that the failure to down-regulate FGF signaling after midbrain-hindbrain boundary specification in RPGRIP1L-deficient conditions may prevent neurogenesis by maintaining early neural cells in their progenitor state.

### FGF/MAPK pathway overactivation manifests as an increased number of SP8-expressing cells and by pMEK1/2 enrichment at the ciliary base

To better characterize the upregulation of FGF signaling, we first investigated in which cells FGF targets were upregulated. We performed immunofluorescence on organoid cryosections for the transcription factor SP8, a known target of FGF8 in the brain [60–62]. In RPGRIP1L KO and in JBTS patient-derived organoids, the number of SP8-expressing cells was increased by 5.7-fold and 4.4-fold, respectively (Fig. 4E). Increased SP8 expression was found mostly, but not only, on polarized progenitor cells of the organoid borders and internal rosettes. We then assessed which cells were producing the FGF8 ligand. Immunostaining analysis on cryosections of day 14 organoids revealed that the strongest FGF8 signal coincided with the apical surfaces of early SOX2^+^/ZO1^+^ apical progenitors. While in control organoids we could only detect FGF8 protein in cells bordering the few internal lumens (Supplementary Fig. 4A), in RPGRIP1L KO and JBTS patient-derived organoids, FGF8 distribution was more widespread. Indeed, FGF8 was enriched on both the apical sides of neural progenitors forming internal lumens (Supplementary Fig. 4A) and the external layer covering the organoid surface (Supplementary Fig. 4B). Thus, the upregulation of FGF signaling observed in transcriptomic experiments may relate both to an increased number of cells expressing the FGF8 ligand and to an increased response of downstream pathway targets such as SP8, also known to activate FGF8 expression [60,61]. To link the FGF pathway upregulation to cilia defects, we analyzed the subcellular localization of pathway members. FGFR1 and FGFR2 and downstream MAPK pathway effectors have recently been reported to be localized at the cilium of epithelial cells in culture [63]. While we could not detect ciliary FGFR localization with available antibodies in our organoids, we found that an activated form of MEK1/2 (pMEK1/2), a MAPK pathway effector, localized at the cilium base in cerebellar organoid cryosections at day 14. More precisely, pMEK1/2 was detected between the centriole labelled with gamma-TUBULIN and the axoneme labelled by ARL13B, thus possibly at the TZ (Fig. 4F). This was the major site of pMEK1/2 localization in non-mitotic cells of organoids from all genotypes analyzed (Supplementary Fig. 4C). In RPGRIP1L KO and JBTS patient-derived organoids, pMEK1/2 localized to the distal end of centrioles even in the absence of an axoneme and showed a global increase in intensity compared to controls, although levels varied between organoids. (Fig. 4F). Thus, the loss of RPGRIP1L increased the activity of the core FGF/MAPK signaling effector pMEK1/2 at the cilium base during cerebellar organoid differentiation, showing FGF pathway overactivation within individual cells.

### RPGRIP1L-deficient organoids display severe neuronal maturation and extracellular matrix defects

To assess whether the impaired neurogenesis observed at early differentiation stages (day 14) prevented the acquisition of a mature neuronal signature in RPGRIP1L-deficient conditions, we analyzed DEG’s profile at later stages. At day 28, we observed a significant downregulation of numerous genes involved in neuronal maturation and function, both in RPGRIP1L KO and JBTS patient conditions (Supplementary Excel Files 4 and 5, respectively) as illustrated by the volcano plots (Supplementary Fig. 5A, B) and on the heatmap representation of normalized RNA expression values over time (Fig. 5A, upper panel). This was confirmed by Single-sample Gene Set Enrichment Analysis (ssGSEA) of neurogenesis and neuronal maturation GO BP terms (Supplementary Fig. 5C).

**FIGURE 5.**
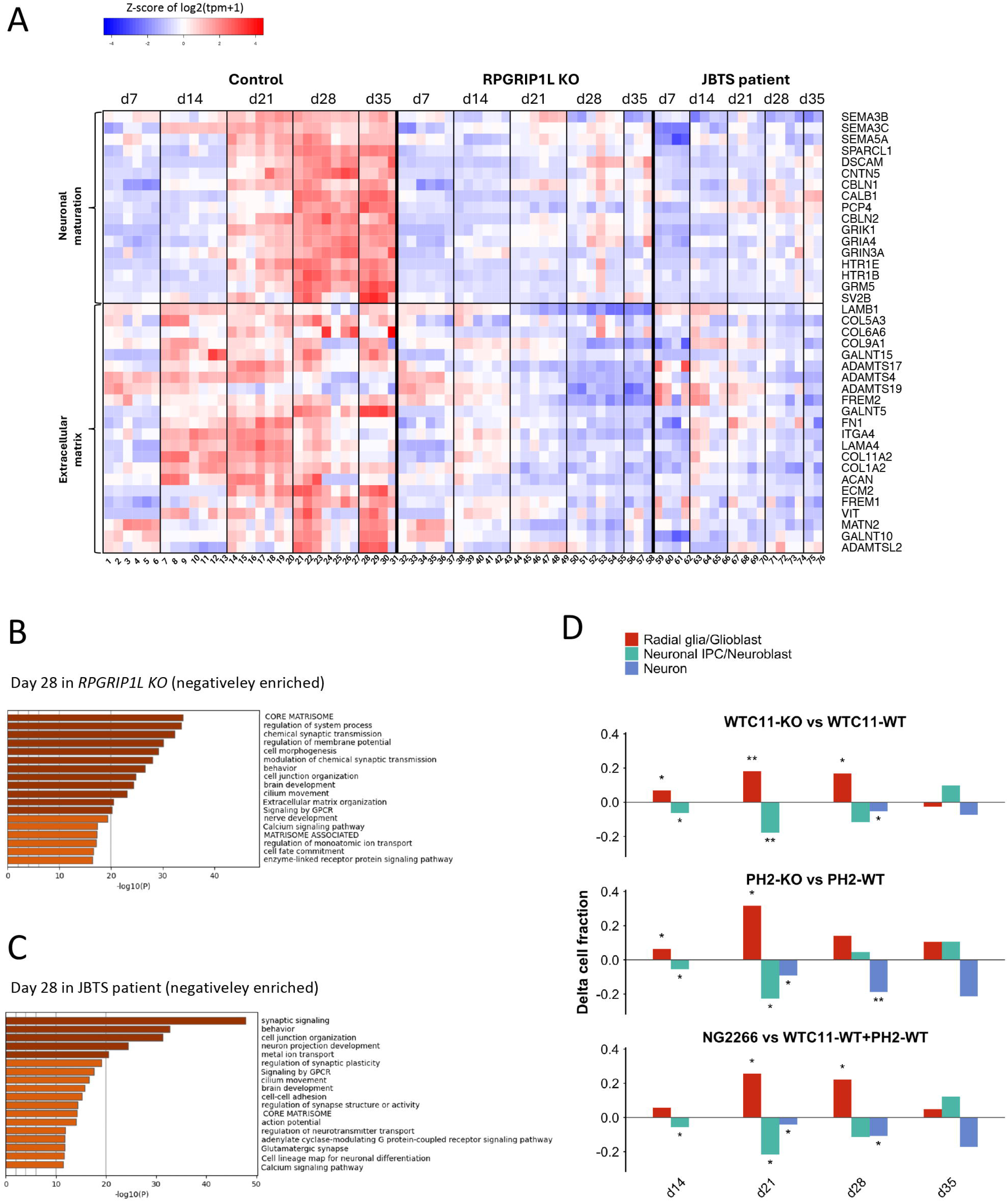
RPGRIP1L-deficient organoids display severe neuronal maturation and extracellular matrix defects as shown by transcriptome analysis (A) Heatmap representing the normalized RNA expression values of selected differentially expressed genes in control and RPGRIP1L-deficient cerebellar organoids over time. 20 organoids were analyzed per condition. See supplementary Table 2 for number of independent experiments and sample numbering. (B, C) Selected GO-terms overrepresented among RPGRIP1L KO (B) and JBTS patient (C) downregulated genes extracted from bulk RNA seq DESeq2 lists at day 28. (D) Graph showing ratios of cell populations in RPGRIP1L- deficient vs control organoids at different stages of differentiation, estimated by bulk deconvolution of scRNAseq data from human first trimester brain atlas^64^. Negative values indicate reduced ratio of the cell type in RPGRIP1L-deficient dataset, while positive values indicate increased ratio.

In absence of functional RPGRIP1L protein we also observed a severe reduction of several genes encoding extracellular-matrix (ECM) components, such as collagens and laminins, as well as integrin receptors (Fig. 5A lower panel; Supplementary Fig. 5A, B).

Gene ontology analyses performed to identify overrepresented gene categories in our DEG lists corroborated these results. Indeed, the most significantly enriched GO terms among the downregulated genes in both RPGRIP1L KO (Fig 5B, Supplementary Excel File 6) and JBTS patient (Fig 5C, Supplementary Excel File 7) conditions at day 28 included brain development, synaptic signaling/chemical synaptic transmission, regulation of membrane potential/action potential, but also core matrisome and extracellular matrix organization.

To refine the comparison between control and RPGRIP1L-deficient organoids by estimating the relative abundance of progenitor and neuronal populations, we performed cell type deconvolution using BayesPrism [64] which predicts fractions of different cell types in bulk RNA-seq samples using as a reference single-cell atlas. In our study we implemented BayesPrism with scRNAseq data from an atlas of the first-trimester fetal human brain [65].

All RGPR1IPL-deficient organoids from both RPGRIP1L-KO hiPSC lines and the JBTS patient line, showed an important increase compared to controls in radial glial (early neural progenitor) cell fraction, peaking at days 14 and 21, at the expense of neuronal intermediate progenitor and neuron fractions (Fig. 5D).

Altogether these results demonstrated a neuronal maturation impairment, following FGF upregulation and neurogenesis defects at early stages, and highlighted a concomitant transcriptional downregulation of major ECM components.

### Blocking FGF signaling rescues neurogenesis and Purkinje cell progenitor markers in RPGRIP1L-deficient cerebellar organoids

To test the hypothesis that the sustained and strong FGF signaling upon RPGRIP1L depletion is at the origin of neurogenesis and neuronal differentiation impairment, we designed a drug rescue experiment. Given that FGF8 expression is crucial for midbrain-hindbrain boundary formation at early stages and that, in RPGRIP1L-deficient organoids, the over-expression of FGF ligands and of their downstream targets started before day 14 and continued until day 21, we treated RPGRIP1L-deficient organoids (both KO and JBTS patient lines) with the pan- FGFR inhibitor BGJ-398 (Infigratinib) [66,67], from day 11 to day 18 (Fig. 6A). Notably, BGJ- 398 treatment restored a normal size of RPGRIP1L-KO and JBTS patient organoids (Fig. 6B).

**FIGURE 6.**
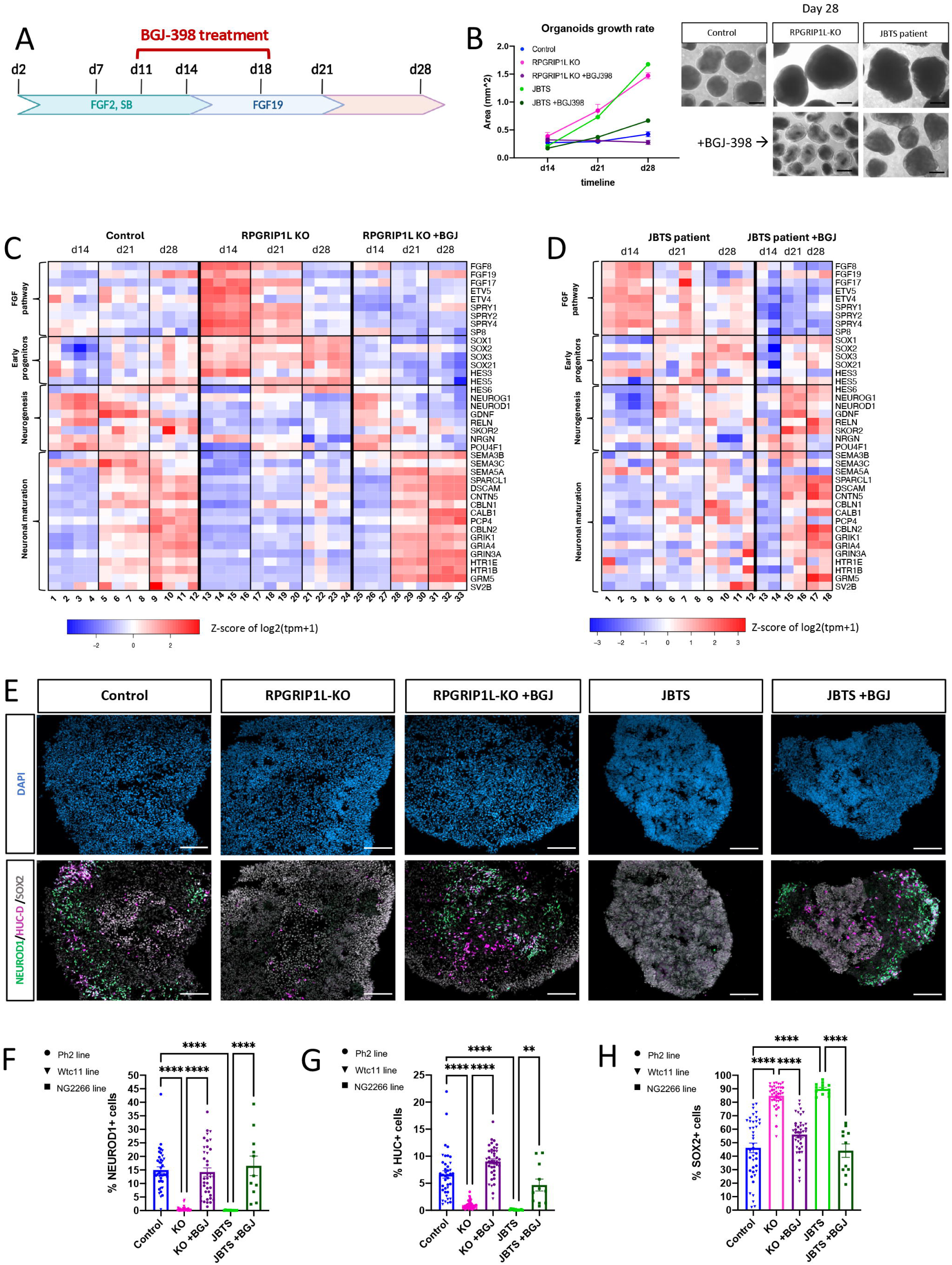
Blocking FGF signaling with BGJ-398 drug rescues organoid size and neurogenesis in RPGRIP1L-deficient cerebellar organoids. (A) Scheme representing the experimental plan for organoid treatment with FGFR-inhibitor (BGJ-398) along the 3D cerebellar differentiation protocol. Bracket on the top represent the time-window of BGJ-398 treatment. (B) Graph representing the growth rate of cerebellar organoids (left) and representative bright field images of cerebellar organoids at day 28 (right). Organoid area was measured every 7 days from day 7 to day 35 of the differentiation protocol. Data are shown as the mean ± SEM of average organoid area from several independent experiments. Control and RPGRIP1L KO: N=3 (Wtc11), JBTS patient: N=1 (NG2266 line); n=10-60 organoids analyzed per experiment per condition. Scale bar, 500 µm. (C, D) Heatmaps representing the normalized RNA expression values over time of selected DEGs in control and RPGRIP1L KO (C) or control and JBTS patient-derived (D) cerebellar organoids, untreated or treated with BGJ-398. For sample numbering and number of independent experiments see Supplementary Table 3 (for C) and Supplementary Table 4 (for D). At least 20 organoids were analyzed per condition per experiment. (E) Immunofluorescence for SOX2 (gray), NEUROD1 (green) and HUC/D (magenta) on cryosections from control and RPGRIP1L-deficient organoids untreated or treated with BGJ- 398 at day 14. Nuclei are stained with DAPI (blue). Scale bar, 100 µm. (F-H) Graphs showing the % of NEUROD1+ (F), HUC/D+ (G) and SOX2+ (H) cells over DAPI+ nuclei., counted in > 15 sections from different organoids per condition per experiment. Controls and RPGRIP1L KO: N=3 (Wtc11 N=2 – triangular dots, Ph2 N=1 – circular dots), JBTS: NG2266 N=1 squared dots. Each dot in the graph represents one organoid; mean ± SEM are shown. Asterisks denote statistical significance according to One-Way ANOVA Kruskal-Wallis test with Dunn’s correction (**P<0.0021; ****P<0.0001). N: number of independent experimental replicates.

We then performed bulk RNAseq analysis on the BGJ treated samples and compared them to their untreated counterparts. Heatmap representation of normalized RNA expression values for selected genes over time clearly show that BGJ treatment was able to rescue the expression of FGF pathway related genes, as well as neurogenesis, neuronal differentiation and extracellular matrix genes in both RPGRIP1L KO (Fig. 6C and Supplementary Fig. 6A) and JBTS patient organoids (Fig. 6D and Supplementary Fig. 6B) from several independent experiments (see Supplementary Excel File 8 for normalized gene expression data).

Quantitative PCR analysis confirmed these results: the RNA expression levels of FGF8 and its downstream targets (ETV4, ETV5, SPRY1) were completely restored by the drug treatment in both RPGRIP1L KO and JBTS patient organoids at all timepoints analyzed (Supplementary Fig. 6C-F). The expression levels of proneural genes NEUROG1 and NEUROD1, as well as Purkinje markers SKOR2 and CALB1, analyzed by quantitative PCR were partially rescued in RPGRIP1L KO and JBTS patient organoids upon BGJ-398 treatment, in particular at their peak of expression (Supplementary Fig. 6G-J). Immunofluorescence analysis on cryosections of day 14 organoids corroborated these results, showing a significant rescue in the number of positive nuclei for either NEUROD1, HUC/D or SOX2 markers, upon BGJ-398 treatment in both RPGRIP1L KO or JBTS patient-derived organoids (Fig. 6E-H). The expression level of the Purkinje progenitor marker OLIG2 was also significantly rescued upon BGJ-398 treatment in both RPGRIP1L KO and JBTS patient-derived organoids, as shown by immunofluorescence analysis at day 21 (Supplementary Fig. 6K).

To confirm that BGJ-398 indeed acted on FGF signaling, we analyzed both targets and pathway members by immunostaining on RPGRIP1L KO and JBTS patient-derived organoids with or without drug treatment. BGJ treatment significantly reduced the number of SP8- expressing cells in both RPGRIP1L-deficient conditions (Fig 7A). In addition, pMEK1/2 amount at cilia base was also significantly reduced by the drug treatment (Fig. 7B and Supplementary Fig. 7A).

**FIGURE 7.**
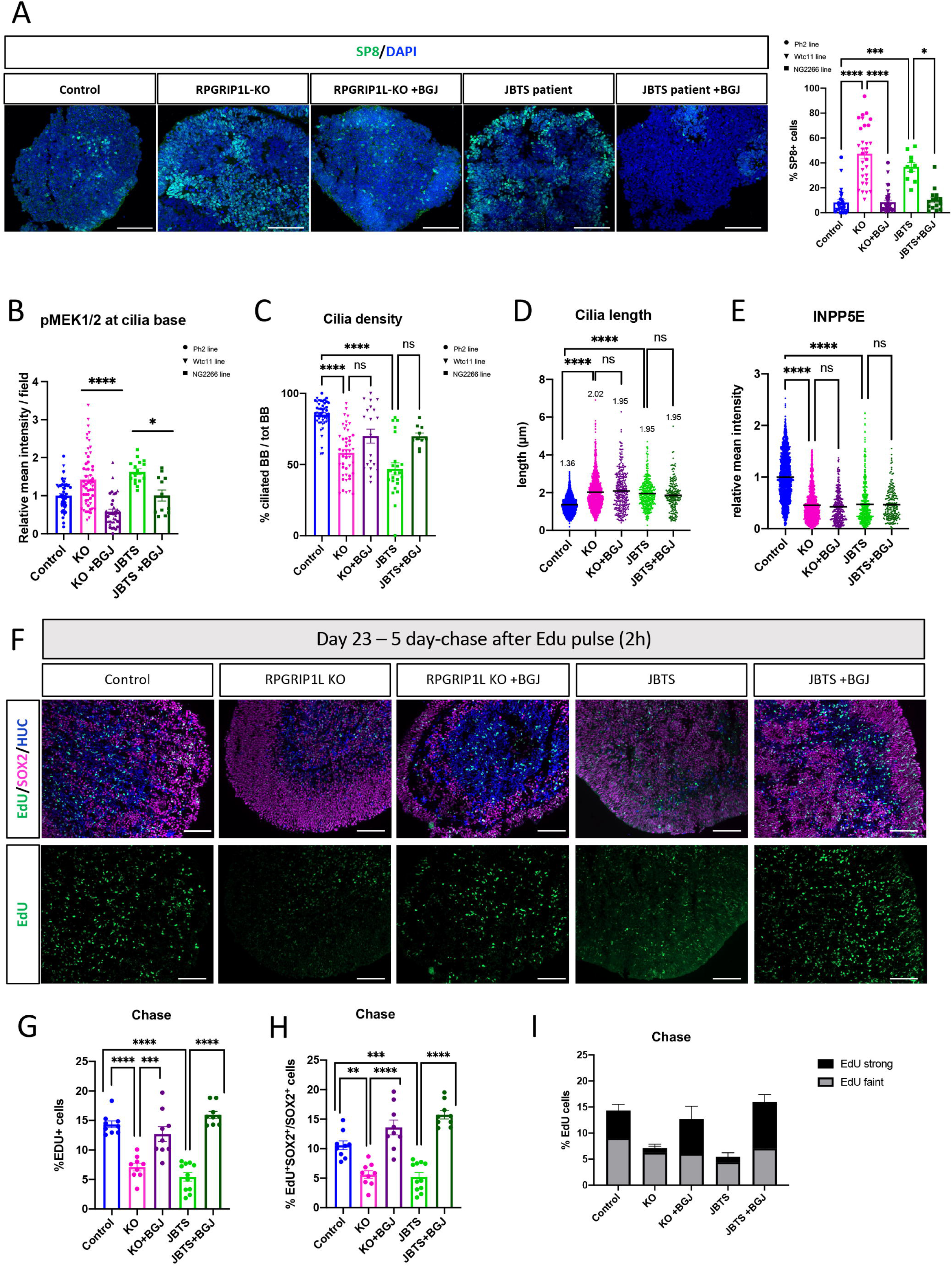
Neural progenitor proliferation is increased in RPGRIP1L-deficient organoids and can be rescued by blocking FGF signaling without restoring cilia features (A) Left: Immunofluorescence for SP8 (green) on cryosections from day 21 control and RPGRIP1L-deficient organoids untreated or treated with BGJ-398. Nuclei are stained with DAPI. Scale bar, 100 µm. Right: Graph showing the % of SP8+ over DAPI+ nuclei, counted in several sections from different organoids per condition. Controls and RPGRIP1L KO: N=3 (Wtc11 N=2 – triangular dots, Ph2 N=1 – circular dots), JBTS: NG2266 N=1 squared dots. Each dot in the graph represents one organoid; mean ± SEM are shown. Asterisks denote statistical significance according to One-Way ANOVA Kruskal-Wallis test with Dunn’s correction (*P<0.033; ***P<0.0002; ****P<0.0001). (B) Graph showing the relative mean intensity of pMEK1/2 labeling adjacent to gamma-Tubulin+ basal bodies, quantified in > 10 sections from different organoids per condition per experiment. Controls and RPGRIP1L KO: N=3 (Wtc11 N=2 – triangular dots, Ph2 N=1 – circular dots), JBTS: NG2266 N=1 squared dots. Each dot in the graph represents the mean of all pMEK1/2 ROIs measured in one field (one organoid) ; mean of the means ± SEM are shown. Asterisks denote statistical significance according to Welch’s t test (***P<0.0002; ****P<0.0001). N: number of independent experimental replicates. (C-E) Graphs showing the quantification of cilia density (C), ciliary length (D) and INPP5E amount in cilia (E) in control and RPGRIP1L-deficient organoids treated or not with BGJ-398. For cilia density, each dot in the graph represents one organoid; mean ± SEM are shown. For cilia length and INPP5E amount, measures for individual cilia (>200) are represented and the mean value is indicated. Asterisks: statistical significance according to One-Way ANOVA Kruskal-Wallis with Dunn’s correction (****P<0.0001). Control and RPGRIP1L KO treated with BGJ: N=2 Wtc11 – triangular dots; JBTS patient: NG2266 N=1 – squared dots. N: number of independent experimental replicates. (F) Immunofluorescence for SOX2 (magenta), HUC/D (blue) and EdU (green) on organoid cryosections at day 23 – 5 days after the 2h EdU pulse (chase experiment), from control and RPGRIP1L-deficient organoids untreated or treated with BGJ-398. (G, H) Graphs showing the % of EdU+ cells over the DAPI+ nuclei (G) and the % of EdU+/SOX2+ cells over the total number of SOX2+ cells (H) at day 23 (chase experiment). Cells were counted in several sections from different organoids per condition. Control and RPGRIP1L KO: N=1 (Wtc11); JBTS: N=1 (NG2266). Each dot in the graph represents one organoid; mean ± SEM are shown. Asterisks: statistical significance according to Ordinary-Way ANOVA after normal distribution testing (**P<0.0021; ****P<0.0001) (I) Graph showing the ratio between “strong” and “faint” EdU labeling in the % of EdU+ cells over total nuclei. Mean ± SEM are shown. N: number of independent experimental replicates. Scale bars: 100 µm.

Finally, to investigate whether reducing FGF signaling could act on cilia formation and content in RPGRIP1L-deficient conditions, we analyzed cilia features in rescue experiments. We found that BGJ-398 treatment had no impact on either cilia density (Fig. 7C), cilia length (Fig. 7D) or ciliary INPPP5E amount (Fig. 7E) in RPGRIP1L KO and in JBTS patient-derived organoids.

Thus, downregulating FGF signaling in RPGRIP1L-deficient cerebellar organoids efficiently rescued the neurogenesis defect without rescuing cilia features.

### Neural progenitor proliferation is increased in RPGRIP1L-deficient organoids and is rescued by blocking FGF signaling

To test whether the size increase observed upon RPGRIP1L impairment could be due to sustained proliferation caused by FGF overexpression, we assessed the proliferative potential of neural progenitors in control, RPGRIP1L KO and JBTS patient-derived organoids by carrying EdU pulse-chase experiments. We provided a 2h EdU pulse to the organoid cultures at day 18, just before the onset of size increase. Some organoids were fixed 2h after the pulse. At this stage (day 18), all RPGRIP1L-deficient organoids showed an increase of EdU positive cells compared to controls (1.6-fold in RPGRIP1L KO, 2.6-fold in JBTS patient) (Supplementary Fig. 7B, C). Of note, similar results were obtained when considering the number of double EdU^+^/SOX2^+^ cells among the population of SOX2^+^ cells, indicating more progenitors in S-phase (Supplementary Fig. 7D). BGJ-398 treatment significantly reduced the number of EdU-incorporating cells in RPGRIP1L KO but not in JBTS patient-derived organoids (Supplementary Fig. 7B-D). This difference may reflect distinct cell cycle dynamics between the two conditions, and/or higher levels of FGF ligand expression in patient-derived organoids (as shown in Fig. 4C) which could delay the response to FGFR inhibition.

A chase assay was performed by removing EdU from the medium after 2 hours and fixing organoids five days after the EdU pulse (i.e. day 23). The number of EdU label-retaining cells was strongly reduced in both RPGRIP1L KO and JBTS patient conditions compared to controls (∼2-fold decrease) (Fig. 7F, G). This indicates that, in the time-window analyzed (between day 18 and day 23), RPGRIP1L KO and JBTS patient-derived cells proliferated more than controls, leading to the dilution of most EdU labeling. BGJ-398 treatment significantly restored the number of label-retaining cells in both RPGRIP1L KO and JBTS patient-derived organoids (Fig. 7F, G). EdU co-labeling with the progenitor marker SOX2 and the quantification of double EdU^+^/SOX2^+^ cells among the SOX2^+^ population showed similar results (Fig. 7H).

In addition, when dividing EdU+ cells into two categories—"strong" and "faint"—based on labeling intensity, we observed that the "strong" EdU+ population, likely representing cells that underwent at most one division after EdU incorporation, was the most affected in both RPGRIP1L-deficient conditions and restored by BGJ-398 treatment (Fig. 7I). Overall, FGF overactivation in RPGRIP1L-deficient cerebellar organoids maintains SOX2+ progenitors in a proliferative state and prevents their switch to neuronal differentiation and maturation, drastically reducing Purkinje cell markers.

## DISCUSSION

A crucial role of primary cilia for proper cerebellar development has been identified in mouse mutants for ciliary genes, which present varying degrees of cerebellar hypoplasia [26–31]. However, the mechanisms linking mutations in ciliary genes to human cerebellar impairment, as well as the developmental origin of specific JBTS patient cerebellar features, remain largely unknown. Importantly, critical aspects of this disease may be missed in mouse models, as cerebellar formation in humans presents unique features [18,19,68], and JBTS genes dysfunction does not necessarily impact ciliogenesis the same way in both species [42]. To overcome this issue, in this study we leveraged human cerebellar organoids to investigate the formation of early cerebellar progenitors under healthy and pathological conditions linked to mutations in RPGRIP1L, a JBTS causal gene. Our study uncovered an increased proliferation of neural progenitors and a reduction of neurogenesis in RPGRIP1L- deficient cerebellar organoids. These defects are correlated with strong upregulation of several FGF pathway related genes and increased levels of phosphorylated MEK1/2 pathway effector at the cilium base. Rescue experiments by blocking FGF receptors (FGFRs) demonstrate the causal link of FGF up-regulation to both neurogenesis-proliferation imbalance and the reduction of Purkinje cell formation.

Given the important role of RPGRIP1L for ciliary functions, we analyzed cilia formation and ciliary content in RPGRIP1L-deficient cerebellar organoids compared to controls. We showed that the RPGRIP1L protein is necessary for correct TZ organization and differentially required for ciliogenesis in early neural progenitors. Indeed, its loss causes a reduced number and an increased length of cilia in non-polarized progenitors, but only rare morphological defects in cilia of polarized progenitors. This difference may result from variations in ciliogenesis pathways and reinforces existing literature indicating a cell type-specific requirement for RPGRIP1L at the ciliary transition zone. Indeed, studies in mice have demonstrated that, in specific cell types, a compensatory mechanism involving the RPGRIP1L paralogue – RPGRIP1 – could overcome RPGRIP1L depletion [53]. Moreover, in depth analysis of cilia distribution and morphology in the mouse cerebral cortex and cerebellum challenged the traditional view that cilia are uniformly present and identical in all brain regions and cell types [69,70,23], a variability that may, at least partially, contribute to the differential vulnerability of brain regions to mutations in ciliary genes. A reduced number of cilia on non-polarized cells could either result from a direct effect of RPGRIP1L loss of function in cilia assembly or stability [45,53] or from an indirect effect via FGF pathway deregulation. Indeed, FGF pathway activation induces cilia lengthening in embryonic tissues and cell lines [71–73] while pathological sustained FGF signalling induces shorter cilia [72]. In addition, a recent paper demonstrated the crucial role of SP8 as a ciliogenesis inducer across embryonic cell types [74]. By performing FGF signaling rescue experiments we found that BGJ-398 drug treatment neither improved nor worsened cilia defects observed in RPGRIP1L-deficient conditions. This strongly suggests that these ciliary alterations arise directly from RPGRIP1L loss of function, while dampening the FGF pathway restores the neurogenesis–proliferation balance downstream of cilia defects.

Our data suggest that defects in RPGRIP1L-deficient cerebellar organoids appear after the initial cerebellum induction stages: while midbrain-hindbrain boundary markers were expressed at day 7 in all genotype conditions, temporal expression analysis of typical cerebellar cell markers revealed a strong reduction of all analyzed markers of the Purkinje lineage, from day 21 to 35. Whether this reflects a defect in Purkinje cell specification and/or in their differentiation remains to be determined. Purkinje cells being early-born neurons in the human cerebellum (8-10 PCW) [17,18], the strong defect in their production in RPGRIP1L-deficient organoids might stem from the general neurogenesis defect. Indeed, the concurrent rescue of neurogenesis and expression of the Purkinje markers OLIG2, SKOR2 and CALB1 following FGFR inhibitor treatment strongly suggests that the two defects are linked. The GCP population appears less affected when considering GCP marker RNA levels. However, we did observe a 50% reduction in the number and staining intensity of BARHL1- expressing cells at day 28, suggesting that the glutamatergic lineage might also be affected at later stages, consistent with the late maturation of GNs in humans [18,49].

A deconvolution analysis of our bulk RNAseq dataset, using the scRNAseq data from a first- trimester fetal human brain atlas [65], confirmed the global maturation delay in RPGRIP1L- deficient organoids compared to controls.

Transcriptome analysis also revealed a severe downregulation of several integrins and ECM components, perturbing the ECM-receptor system in RPGRIP1L-deficient organoids. Although the causes of this alteration are unclear, it might participate in the neurogenesis and neuronal maturation impairment in RPGRIP1L-deficient organoids. Indeed, the expression of ECM molecules is dynamically regulated during brain development and plays prominent roles in neurogenesis, cell migration, neurite outgrowth and synaptogenesis [75–77]. The downregulation of ECM genes may be responsible at least in part for the observed changes in the organization of progenitor zones, with more progenitors lying at the external border of the organoids in RPGRIP1L-deficient embryos than in controls. These changes may also contribute to the neurogenesis phenotype by increasing the exposure of progenitors to external cues present in the medium, such as FGF2.

FGF8 is known as the key morphogen to establish early cerebellar territory in vivo [78]. FGF8 is initially expressed in the isthmic region, the prospective cerebellar vermis, and is required to induce cerebellar formation [79,80]. Notably, the dose, the precise location and the time window of FGF8 expression need to be tightly controlled to allow proper cerebellar development [81]. Ectopically upregulating Fgf8 in cerebellar precursors increases their proliferation and overrides their normal program of cell specification [82]. Moreover, conditional inactivation in the mouse isthmus of Sprouty genes (Spry1-2-4), which are FGF direct targets acting as feedback loop negative regulators [83], results in overactivation of FGF signaling and aberrant cerebellar development [84]. A growing literature emphasizes the role of the FGF pathway in regulating the proliferative/neurogenic balance in early neural development, by promoting progenitor proliferation at the expense of neuronal differentiation during embryonic spinal cord elongation [85–87] and corticogenesis [88,89].

In our cerebellar differentiation protocol, control organoids downregulated FGF8 after day 7, while RPGRIP1L-deficient organoids displayed strong FGF8 up-regulation from day 14 to 21. Consistent with current literature, we showed that the increased levels of FGF ligands (FGF8, FGF17, FGF19) and pathway activation, as indicated by the expression of ETV4-5 target genes, were responsible for the impaired neurogenesis and neuronal differentiation observed upon RPGRIP1L depletion.

We thus addressed the mechanisms that could underlie FGF upregulation upon RPGRIP1L- deficiency. A recent paper showed that, in mouse embryos and in cell culture, FGF receptors 1 and 2 localize to the cilium [63], and that downstream activated MAPK effectors are present at the cilium base. We found that in our cerebellar organoids, the activated phosphorylated form of the MAPK kinase MEK1/2 (pMEK1/2) localized almost exclusively to the cilium base in most cells, except in mitotic cells where it localized to the spindle poles. In RPGRIP1L-deficient organoids, pMEK1/2 was also localized to the basal body, even in cells with no visible cilia. Moreover, the amount of pMEK1/2 at cilia was increased, suggesting that defects in TZ organization may, directly or indirectly, impact pMek1/2 amount at the cilium base. Thus, our data show that 1) FGF signaling is activated at the cilium base in cerebellar progenitors, 2) RPGRIP1L and cilia are not required for FGF signaling activation in these cells, nor for pMEK1/2 recruitment adjacent to basal bodies and 3) on the contrary, RPGRIP1L deficiency leads to a defect in signal attenuation.

In which cell types is FGF signaling active? We revealed the presence of FGF8 protein mainly in apical neural progenitors, suggesting that they are the FGF8 producing cells. In terms of responding cells, both apical and non-apical neural progenitors activate the pathway, as seen with pMEK1/2 and SP8 levels. We thus propose that the FGF pathway activation at the cilium base initiates a positive loop by increasing the number of proliferating progenitors that themselves activate the FGF pathway. We provide evidence that, once engaged, this mechanism could be self-sustaining. We propose that the SP8 transcription factor, a well- known inducer and target of FGF8 [90,91,60], participates in this self-amplification loop, being significantly upregulated at both day 14 and 21 (see DEGs tables) and expressed by an increased number of cells in all RPGRIP1L-deficient conditions.

An essential question raised by our study is how the increased proliferation observed in cerebellar organoids may relate to the selective vermis hypoplasia characteristic of JBTS patients, whose cerebellar hemispheres are comparatively preserved. To the best of our knowledge, there is no specific gene expression marker to discriminate between developing vermis and hemispheres, and organoid models do not recapitulate the spatial complexity of a developing cerebellum. Moreover, we cannot exclude that the context of organoids could favor specific manifestations of the RPGRIP1L mutant phenotype that are not revealed in vivo. Despite these limitations, several hypotheses could explain the apparently opposite phenotype between RPGRIP1L-deficient organoids and JBTS pathology. First, an early over- proliferation could trigger later cell death, as observed upon loss of function of the kinase DYRK1A [92]. Alternatively, an increase in early progenitor numbers could preferentially populate the cerebellar hemispheres at the expense of the vermis. Indeed, lineage studies in mice highlight different medio-lateral distribution of cerebellar VZ clones according to their birth dates [93,94]. Finally, aberrant FGF8 signaling may perturb morphogenetic processes at the midbrain–hindbrain boundary. Alterations in the size and organization of the isthmic organizer have been shown to impair midline fusion of the bilateral cerebellar primordia, a process required for vermis formation [95].

The long-term consequences of impaired differentiation and sustained proliferation in cerebellar organoids will have to be investigated, but it is worth noticing that this phenotype is consistent with data obtained in mouse mutants for two JBTS causal genes, KIF17 [96] and SUFU [97–99]. Interestingly, SUFU variants in humans are associated with JBTS vermis hypoplasia [100, 101], but also represent an increased risk factor for severe infantile medulloblastoma [102,103]. This, together with our data on RPGRIP1L deficiency in cerebellar organoids, raises the puzzling question of potential common mechanisms between early cerebellar progenitor hyperproliferation and later vermis hypoplasia.

Although our model cannot fully address all these questions, the relevance of our study for JBTS pathology is emphasized by the similar phenotypes observed in RPGRIP1L KO organoids and organoids derived from a JBTS patient’s iPSC line. The JBTS patient donor of the NG2266 cell line used in this study, exhibits a very severe neurological phenotype, correlated with vermis hypoplasia and partial aplasia, as revealed by encephalic magnetic resonance imaging (MRI). The patient carries two biallelic heterozygous mutations in the RPGRIP1L gene: a nonsense mutation at the end of the first C2 protein domain, and a splice variant that results in the generation of a stop codon at the beginning of the second C2 domain. Given the importance of both C2 domains for RPGRIP1L function [104,105,26,106,43], a stop codon in these domains is likely to impair protein functionality. Indeed, similar to the RPGRIP1L KO condition, JBTS patient (NG2266)-derived organoids lacked RPGRIP1L protein at the ciliary transition zone. This further underscores the strong phenotypical similarity observed between RPGRIP1L KO and JBTS patient (NG2266)-derived organoids. Furthermore, targeted pharmacological inhibition of FGFR using BGJ-398/Infigratinib, an FDA-approved drug currently used in oncology and to treat hypochondroplasia [67], is sufficient to rescue the proliferative-neurogenic balance and to restore Purkinje cell lineage formation, both in RPGRIP1L knockout hiPSC lines and in the Joubert syndrome patient line, establishing a mechanistic link between FGF overactivation and the observed phenotypical alterations.

More generally, it will be crucial to determine whether the molecular alterations reported in this study have broader relevance by also occurring in cerebellar organoids with mutations in other JBTS genes, or if they are specific to RPGRIP1L depletion. It will also be particularly helpful to investigate whether FGF signaling could be specifically disrupted in cerebellar organoids from severely affected patients, as it might be of predictive value for the severity of the neurological phenotype.

In conclusion, with this work we provide an example of a cerebellar organoid approach to investigate potential pathogenetic mechanisms underlying a human disease. Our results uncover neurogenesis defects driven by increased FGF signaling, thereby offering a novel framework for investigating the etiology of cerebellar defects in JBTS and other neurodevelopmental diseases. We further highlight that RPGRIP1L ciliary protein performs complex rheostat functions in signaling, revealing its crucial role for proper FGF signal attenuation in human neural progenitors, while in other cellular contexts in mice, Rpgrip1l is necessary to reach the highest level of Hedgehog activation [46].

## MATERIAL AND METHODS

### Ethics

The work performed in this manuscript complies with all relevant ethical regulations. The obtaining and manipulation of human iPSC lines were performed under ethical agreements from the French Ministry of Higher Education and Research DC-2025-7003 (for the UCSFi001-A and PCIi033-A lines, control and RPGRIP1L-KO) and DC-2025-7496 (for the JBTS patient line) and under GMO authorization number DUO-9509. Formal consent was obtained from the patients (for adult donors) or their parents (for child’s donors).

### Human iPSC production, culture and quality control

Control and RPGRIP1L mutant hiPSC clones were generated by CRISPR/Cas9-mediated genome editing as described [42], in two different parental hiPSC lines: PCiI033-A (PHENOCELL (PLI)) product reference PCi-CAU2) and UCSFi001-A (deposited at the Coriell Institute for Medical Research under the identifier GM25256). Genome editing was performed at passage 44 for UCSFi001-A (Wtc11) and at passage 41 for PCIi033-A (Ph2). The patient-derived iPSC line NG2266 was derived as described [51]. For all clones, genomic stability was assessed by detection of recurrent genetic abnormalities using the iCS-digital PSC test, provided as a service by Stem Genomics (https://stemgenomics.com). The pluripotency of each iPSC clone was confirmed using the STEMdiff Trilineage Differentiation kit (STEMCELL technologies) according to the manufacturer’s instructions. Pluripotency and lineage differentiation were tested by qPCR using marker genes for iPSCs (OCT4, NANOG), mesoderm (MIXL1), endoderm (SOX17, EOMES) and ectoderm (PAX6, LHX2). hiPSCs were thawed in mTeSR+ medium (Stem Cell Technologies #100-0276) in the presence of Rock- inhibitor Y-27632 (5μM, Stemgent-Ozyme #04-0012) and cultured on Matrigel (Corning, VWR #354277) coated plates under standard conditions at 37°C upon confluency of 70-80%. The medium was changed every two days. Passaging was performed using ReLeSR (Stem Cell Technologies #05872) and testing for potential mycoplasma contamination was performed regularly by Eurofins genomic (Mycoplasmacheck). Accutase (Stem Cell Technologies #07920) was used for the dissociation of hiPSC colonies into single cells prior to the differentiation protocol.

Clinical features of the patient COR201 and characterization of the mutant Identification of the RPGRIP1L RNA products derived from the NG2266 hiPSC c.2304+1G>T splice variant allele.

The NG2266 cell line is derived from the male patient COR201 [51] who was last examined at the age of 17. He presented a molar tooth sign on brain MRI and his neurological examination revealed hypotonia, psychomotor delay, neonatal breathing dysregulation, ocular-motor apraxia, nystagmus and severe intellectual disability. Post-axial polydactyly of the left hand was present at birth. During evolution, he suffered severe visual impairment and strabismus, severe renal dysfunction (chronic renal failure at the age of 11, which required renal transplantation at age 14), as well as chronic liver disease.

We demonstrated by sequencing PCR products from NG2266 hiPSCs cDNA that the splice variant c.2304+1G>T resulted in intron retention leading to an early stop codon at position 788 (S788X). Primer 1 (Forward: GTATCCATAAACTAGAAGCCCAAT) was designed at the junction between exon 14 and 15; primer 2 (Reverse: GGGTTAAGGTTACAGGGGATCTC) was placed in the intron between exon 16 and 17; Primer 3 (Reverse: AATATTCCTGAGATACACCTGTC) was positioned at the junction between exon 17 and 18 (Supplementary Fig. 8A). PCR products were analyzed on agarose gel and showed that primer pair 1-2 amplified an 813 bp region in the cDNA derived from JBTS patient (NG2266) hiPSCs but not from the control condition (Supplementary Fig. 8B). Primer pair 1-3 amplified a 1009 bp region, both in the control and in JBTS patient cDNA. As expected, NG2266 second allele (c.2050C>T/p.Q684X) presents a nonsense mutation that does not affect the cDNA structure, resulting in a PCR product of the same size as control alleles. On the contrary, using the primer pair 1-3, the c.2304+1G>T allele could not be amplified from cDNA given that intron retention would give rise to a 3969 bp PCR product (Supplementary Fig. 8B’).

Primer pair 1-2 PCR product from NG2266 hiPSCs was sequenced to confirm the intron retention. An ATG stop codon appears 29 nt after the beginning of the intron sequence, most likely giving rise to a truncated form of the protein (Supplementary Fig. 8C). Moreover, an additional mutation was identified in the exon 16: c.2231 G>A (p. R744Q).

### Differentiation of hiPSCs into cerebellar organoids

For organoid production, iPSC lines were thawed at Passage 7-11 after genome editing, so P44 + [7–11] for WTC11 and P41 + [7–11] for Ph2. NG2266 JBTS patient hiPSC line was thawed at passage 16-20 (after the loss of SeV vectors). After two amplification passages, hiPSC lines (characterized in [51,53]) were dissociated into single cells using Accutase (Stem Cell Technologies #07920) and resuspended in mTeSR+ medium supplemented with 10 μM ROCK inhibitor. Then, cells were promptly re-aggregated using AggreWel 800 microwell plates (StemCell Technologies) following the manufacturer’s instructions. Cells were seeded at a density of 1.8 × 10^6 cells per well (6,000 cells per microwell). This corresponded to day 0 of the cerebellar differentiation protocol. Two days later the entire medium was replaced with the chemically defined medium (gfCDM), slightly modified compared to [33], consisting of vol:vol Isocove’s modified Dulbecco’s medium (Thermo Fisher #12440053)/Advanced DMEM/F-12 (Thermo Fisher #12634028), chemically defined lipid concentrate (1% v/v, Thermo Fisher #11905031), monothioglycerol (450 μM, Sigma #M6154), apo-transferrin (15 μg/ml, Sigma #T1147), crystallization-purified BSA (5 mg/ml, >99%, Sigma #5470), and 50 U/ml penicillin/50 μg/ml streptomycin (P/S, Thermo Fisher #15140122). The medium was also supplemented with insulin (7 μg/ml, Sigma #I9278) and used from day 2 to day 21. Different factors were added to the medium at different timepoints as represented in Fig. 1A. Recombinant human basic FGF (FGF2, 50 ng/ml, PeproTech # 100-18B) and SB431542 (SB, 10 μM, Stemgent-Ozyme #04-0010) were added to the culture on day 2. On day 4, floating aggregates were transferred from microwell plates to ultra-low attachment dishes (Corning #3261) and total medium change with drug refreshment was performed. On day 7 embryoid bodies(EBs)-containing dishes were transferred to an orbital shaker (3 mm orbit, Thermo Scientific #88881102B) and maintained under orbital agitation of 100 rpm until the end of the protocol. Total medium change with 2/3 of the initial amount of FGF2 and SB431542 was performed on day 7, day 9 and day 11. On day 14, FGF2 and SB431542 were replaced by medium change supplemented with recombinant FGF19 (100 ng/ml; PeproTech #100-32). The same medium change was also performed at day 16 and 18. From day 21 onward, the aggregates were cultured in Neurobasal medium (Thermo Fisher #21103049) supplemented with GlutaMax I (Thermo Fisher #35050061), N2 supplement (Thermo Fisher #17502048), and P/S 1% (Thermo Fisher #15140122). Total medium changes were performed on day 23, 25, 28, 30 and 32. Recombinant human SDF1 (300 ng/ml, PeproTech #300-28A) was added to the culture from day 28 onward (Fig. 1A).

For pan-FGFR inhibition, BGJ-398/Infigratinib [66,67] was first added at day 11 at a concentration of 100 nM and renewed on days 14 and 16. At day 18 total medium was replaced by fresh medium devoid of BGJ-398.

### EdU treatment and labeling

Cerebellar organoids at day 18 were incubated using 10 μM EdU (Click-iT EdU Imaging Kit, Invitrogen) dissolved in growth medium for 2 h at 37 °C and then washed twice with PBS. Around 30 organoids per genotype were collected and fixed for 20 minutes at room temperature in 4% cold PFA. The remaining organoids, around 30 per genotype, were kept in culture after washing for a 5-day chase until day 23 of the differentiation protocol, when they were collected and fixed in the same way.

Fixed organoids were embedded in OCT following the procedure described in the section Organoids embedding and Cryosectioning. EdU labeling detection on cryosections was performed according to the Click-iT EdU Imaging Kit manufacturer’s instruction at the end of our immunostaining protocol (refer to the following section “Immunofluorescence”).

### Organoid Size Analysis

To monitor the organoid size, bright-field images were acquired at different time-points (every seven days). Organoid area was calculated by manually dragging a line with the freehand selections tool on Fiji (ImageJ; National Institutes of Health) and subsequently measuring the object area. Organoid area was measured in several independent experiments (N=3 Ph2 line, N=3 Wtc11 line, N=3 NG2266 line), n=10-60 organoids were analyzed per experiment per condition. Results are presented as the mean ± SEM of organoid averages from each individual experiment.

### Organoids embedding and Cryosectioning

Organoids were collected every seven days during differentiation from day 7 to 35, rinsed with PBS and fixed in cold PFA (4 %) for 20 min at 4 °C. Then organoids were rinsed in PBS and incubated in 30 % sucrose in PBS until completely saturated. Cryoprotected organoids were embedded in OCT matrix (Cell Path #KMA-0100-00A) in plastic molds and stored at -80°C. 12 μM cryostat sections were prepared and stored at -20 °C.

### Immunofluorescence

Organoids were collected at different time points during differentiation, rinsed with PBS and fixed in cold PFA (4%) for 20 min at 4 °C. Organoids were then washed in PBS and incubated in 30% sucrose in PBS until completely saturated. Cryoprotected organoids were embedded in OCT embedding matrix (Cell Path #KMA-0100-00A) and stored at -80 °C. 12 μm cryostat sections were prepared and stored at -80 °C. Cryostat sections of organoids were washed in PBS/0.1% Triton X-100. An optional step of antigen-retrieval with Citrate-Buffer was performed only for cilia staining (INPP5E and Gamma-TUBULIN antibodies). After washing, cryosections were incubated with a blocking solution containing 10% NGS in PBS/0.1% Triton-X100 for at least 2 hours. The sections were incubated overnight at 4 °C with primary antibodies (Supplementary Table 5) diluted in a solution containing 1% NGS in PBS/0.1% Triton-X100. Next day, the slides were washed 3 x 10 min with PBS/0.1% Triton-X100 and incubated with secondary antibodies at 1:400 dilution (Supplementary Table 6) and DAPI (Merck #1.24653) in 1% NGS/PBS/0.1 % Triton X-100 for 2 hours. After final washing steps with PBS/0.1 %Triton X-100 for at least 1 hour, sections were mounted with Mowiol (Roth #0713.2).

### Image acquisition

Fluorescence image acquisition of cerebellar progenitors and neuronal markers was performed at room temperature using a Zeiss Observer Z1 equipped with an Apotome module, an Axiocam 506 monochrome cooled CCD camera and a 20x objective. Z-stacks with 1 μM step were acquired. All images were processed with Zen software (blue edition).

For cilia analyses, Z-stacks images with 250 nM steps were acquired using a confocal microscope (LSM 980 upright, Carl Zeiss AG) and a 63x oil objective with a NA 1.4. Phospho- MEK1/2 images used for quantification were acquired as Z-stacks with 250 nM steps with LSM980 confocal microscope using a 63x oil objective with a NA 1.4 and a zoom of 2. All confocal images were acquired using a monochrome charge-coupled device camera and processed with Zen software (black edition).

For Super-resolution imaging of the pMEK1/2 signal, the LSM980 Airyscan confocal (Zeiss) was used in SR mode and image stacks were acquired every 20 nM with a 63x objective and a zoom value of 3.

### Quantification of immunofluorescence staining and statistical analyses

The number of positive cells in immunofluorescence staining was quantified on 7µm Z- projections using the Cellpose environment (https://cellpose.readthedocs.io/en/latest/installation.html) for cell segmentation. Each image was segmented using the DAPI channel to generate a mask identifying all nuclei. The intensity of the target channel was then measured for each nucleus by uploading the MAX- projection image and the corresponding ROI mask (generated with Cellpose) into Fiji (ImageJ; National Institutes of Health). Mean intensity was calculated for each ROI, and a threshold was applied to distinguish marker-positive cells from background immunostaining. Only cells above this threshold were counted, and the ratio to the total number of nuclei (ROIs in the mask) was determined. Nuclei smaller than 10 µm² were excluded, as they were considered pycnotic cells.

For quantitative analyses of ZO1 staining at the apical side of the cells, positive areas were defined and measured by manually setting common thresholds in Fiji for all genotypes. For each image, the DAPI positive area was also determined and measured by manually setting thresholds. The percentage of ZO1 positive area per organoid slice was determined after dividing the ZO1 calculated area by the DAPI area.

Analysis of ciliary density was performed by manually counting the number of cilia axonemes (INPP5E staining) over the number of basal bodies (Gamma-Tubulin staining). Measurements of axonemal INPP5E intensity and ciliary length were performed on unprocessed images (raw data) using the segmented line tool on Fiji to draw ROIs corresponding to each cilium. Mean gray value and length were then extracted for each ROI of an image. INPP5E mean intensity is calculated by subtracting the mean background value of an image to the mean gray value of each ROI in the image. Data are represented as “relative mean intensity” by setting the mean value of all control cilia (ROIs) at 1. After quantifications were performed, representative images were processed by means of background subtraction and contrast settings via Fiji.

pMEK1/2 quantifications were performed on Z-stacks (7.5µ thickness), tracing ROI manually at the base of each cilium between the basal body stained with GAMMA-TUBULIN and the axoneme labelled by ARL13B. The mean intensity of each ROI was measured using FIJI software, after subtracting the background value. The background (cytoplasmic pMek1/2) intensity value was obtained by measuring fluorescence intensity in two ROIs (of the same surface area as the one at the base of the cilium) adjacent to the cilium base and calculating their mean intensity. For each field of 67.3µm X 67.3µm, 20 to 100 ROIs were measured and the mean value for each field was plotted.

All data are presented as mean ± SEM. One-Way ANOVA Kruskal-Wallis and Dunn’s test or Welch’s t test were performed as indicated in the figure legends and the following statistical significances were considered: *P<0.033, **P<0.0021, ***P<0.0002 and ****P<0.0001. All statistical data analysis and graph illustrations were performed using GraphPad Prism (GraphPad Software).

### Total RNA isolation and qPCR analyses

Control, RPGRIP1L KO and JBTS patient-derived cerebellar organoids from different hiPSCs lines (Ph2, Wtc11, NG2266) and several independent experiments were collected every seven days from day 7 to 35 of the differentiation protocol and washed with PBS. RNA was isolated using the RNeasy Kit (Qiagen #74104) and RNAse-free DNase Set (Qiagen #79254) following manufacturer’s instructions. 1µg of isolated RNA was retro-transcribed into cDNA using the GoScript Reverse Transcriptase Kit (Promega #A5001). For quantitative real-time PCR, a cDNA dilution (equal to 15 ng of starting RNA per well) was used in a Maxima SYBR Green/ROX qPCR Master Mix 2x (Thermo Scientific #K0222). Duplicate reactions for each sample were prepared and the samples were run in a Step One Real-Time PCR System Thermal Cycling Block (Applied Biosystems #4376357). GAPDH housekeeping gene was used as reference. Primer pairs are listed in Supplementary Table 7. The analysis of real-time data was performed using the included StepOne Software version 2.0 and graphs were generated using GraphPad Prism (GraphPad Software).

### Total RNA isolation and Bulk RNAseq analysis on cerebellar organoids

RNA was isolated as described above using the RNeasy Kit (Qiagen #74104) and RNAse-free DNase Set (Qiagen #79254) from control, RPGRIP1L KO and JBTS patient-derived cerebellar organoids at different timepoint and from several independent experiments (see Supplementary Tables 2-4 for sample details and N). RNA quality was determined by measuring the RNA integrity number (RIN) via the High Sensitive RNA Screen Tape Analysis kit (Agilent Technologies #5067) on the TapeStation system (Agilent Technologies). A RIN above 9.5 was considered as good quality RNA and 250 ng RNA in a total volume of 25 μl was prepared per sample for further procedure. Bulk RNAseq was realized by the Genotyping and Sequencing Core Facility of the Paris Brain Institute (iGenSeq, ICM Paris). Poly A stranded cDNA libraries were built with the kit “Illumina Stranded mRNA Prep” and their quality checked on “AGILENT Tapestation 2100”. 125 ng of each library was sequenced on both strands to produce between 33-44 Millions reads (100 bases long) using a Novaseq6000 ILLUMINA.

RNAseq data processing was performed in Galaxy under supervision of the ARTBio platform (IBPS, Paris). Paired-end RNA reads were aligned against the homo sapiens hg38 genome by using the STAR aligner (v2.7.10, Galaxy wrapper v4) [107] and the gene annotations gtf files GRCh38.109. Quality of sequencing was controlled with fastQC (Galaxy wrapper v0.74+galaxy0) and MultiQC (Galaxy wrapper v1.11+galaxy1) [108]. Gene expression was assessed by counting sequencing reads aligned to gene exons with featureCounts (Galaxy wrapper v2.0.3+galaxy2) [109]. Raw counts were further converted to normalized gene expression values using the log2(tpm+1) transformation where tpm is the count of transcript aligned reads per length of transcript (kb) per million of total mapped reads (Galaxy tool "cpm_tpm_rpk" v0.5.2).

To construct the diffusion map, the data was first filtered to remove Y_RNA and rRNA genes. Expressed genes were defined as those with log2(TPM+1) > 3 in at least 8 samples (equivalent to about 10% of the dataset). Principal component analysis (PCA) was performed on the remaining 13,160 genes. The first 12 principal components that each explain more than 1% of the total variance were used to construct a diffusion map using the DiffusionMap function from the destiny package (v3.20) in R, with parameters set to k=9 nearest neighbors and 10 eigenvectors [110].

Heatmap visualizations were produced with the Galaxy tool "Heatmap2" (v3.3.0+galaxy0) using the log2(TPM + 1) normalized gene expression values, with row-wise z-score scaling applied before visualization. Differentially expressed genes (DEGs) were selected from the gene raw read counts using DESeq2 (Galaxy wrapper v2.11.40.8+galaxy0) [111] and the Benjamini-Hochberg p-adjusted cutoff 0.05. DESeq2 statistical tables were used for generation of Volcano Plots (Galaxy tool id "volcanoplot" v0.0.5) [112] and for the analysis of overrepresented gene ontology (GO) categories (Metascape software, https://metascape.org/gp/index.html#/main/step1) [113].

Single-sample gene set enrichment analysis (ssGSEA) was performed using the gsva function from the GSVA package (v2.0.7) in R (https://pmc.ncbi.nlm.nih.gov/articles/PMC3618321/). Gene sets were downloaded from the Molecular Signatures Database (MSigDB) (https://www.gsea-msigdb.org/gsea/msigdb).

### Bulk deconvolution

Cell type deconvolution was performed with BayesPrism [64] using a first- trimester fetal human brain atlas [65] as the single-cell reference. Among the 1.6m cells in the atlas, we selected only the cells sequenced using the Chromium (10X Genomics) single- cell reagent kit version 3 to avoid bias in the technology version. We then selected only cells coming from the embryonic phase (<8pcw), and removed all non-neural cells as well as those with the ambiguous "Forebrain" annotation. Oligo and neural crest cells were also excluded due to low cell numbers (<100 cells per cell type). This resulted in a subset of 370,750 cells. Deconvolution was then performed with 3 cell classes “radial glia/glioblast”, “neuronal IPC/neuroblast”, and “neuron”. At each time point, delta values were calculated as the difference in the mean cell fractions between the KO/patient and WT while excluding cell fractions lower than 5%. To test the significance of the changes, the fraction values were first logit-transformed using the logit function from the R package car (v3.1-5) with an adjustment factor of 0.001, then the Student’s t-test was applied. P-values were adjusted for multiple testing using Benjamini-Hochberg correction. Asterisks (*) indicate adjusted p < 0.05.

## ACKNOWLEDGEMENTS

We particularly thank Chloé Chaumeton and France Lam from the IBPS imaging platform I2PS for their help on image acquisition. We thank the IBPS Bioinformatics facility of which C.A. is a former member. We thank the iGenSeq sequencing platform at the ICM for reliable and fast processing of our samples. We are very grateful to Stéphane Nedelec for providing advice on BGJ-398/Infigratinib drug usage in hiPSCs and to Aline Stedman for scientific discussions and advice on manuscript organization. We thank Alexis Eschstruth for his kind contribution to the PCR analysis to identify the cDNA product of JBTS patient c.2304+1G>T splice variant.

---

**Supplementary Figure 1.**
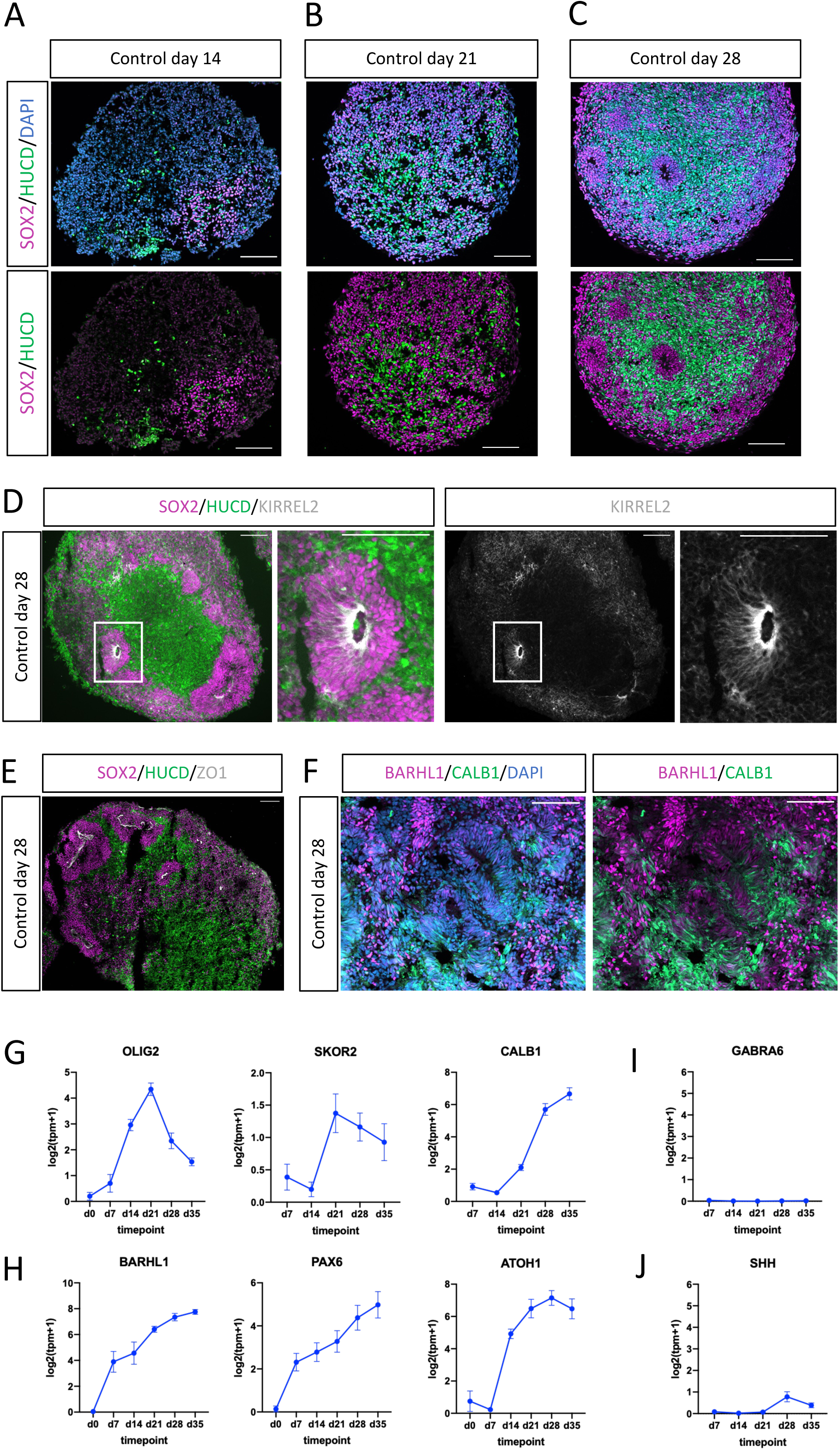
Cerebellar organoids generated from control human induced pluripotent stem cells recapitulate early steps of cerebellar development **(A-C)** Immunofluorescence for SOX2 (magenta, neural progenitors) and HUC/D (green, early- born neurons) on cryosections from control organoids at indicated time-points. Nuclei are stained with DAPI (blue). **(D-F)** Immunofluorescence on cryosections from day 28 control organoids, for KIRREL2 (white), SOX2 (magenta) and HUC/D (green) (D); ZO1 (white), SOX2 (magenta) and HuC/D (green) (E); and BARHL1 (magenta) and CALB1 (green) (F). Scale bars: 100 µm in A-F. **(G-J)** Graph showing normalized temporal gene expression analysis from bulk RNAseq data : Purkinje cell markers: OLIG2, SKOR2, CALB1 (G), Glutamatergic lineage markers BARHL1, PAX6, ATOH1 (H), Granule Neuron marker (*GABRA6*) (I), and *SHH* (J). Data are displayed as mean ± SEM and were obtained from both Ph2 and Wtc11 cell lines. Day 0 (hiPSCs): N=2; day 7-35 organoids: see Supplementary Table 2 for number of independent experimental replicates.

**Supplementary Figure 2.**
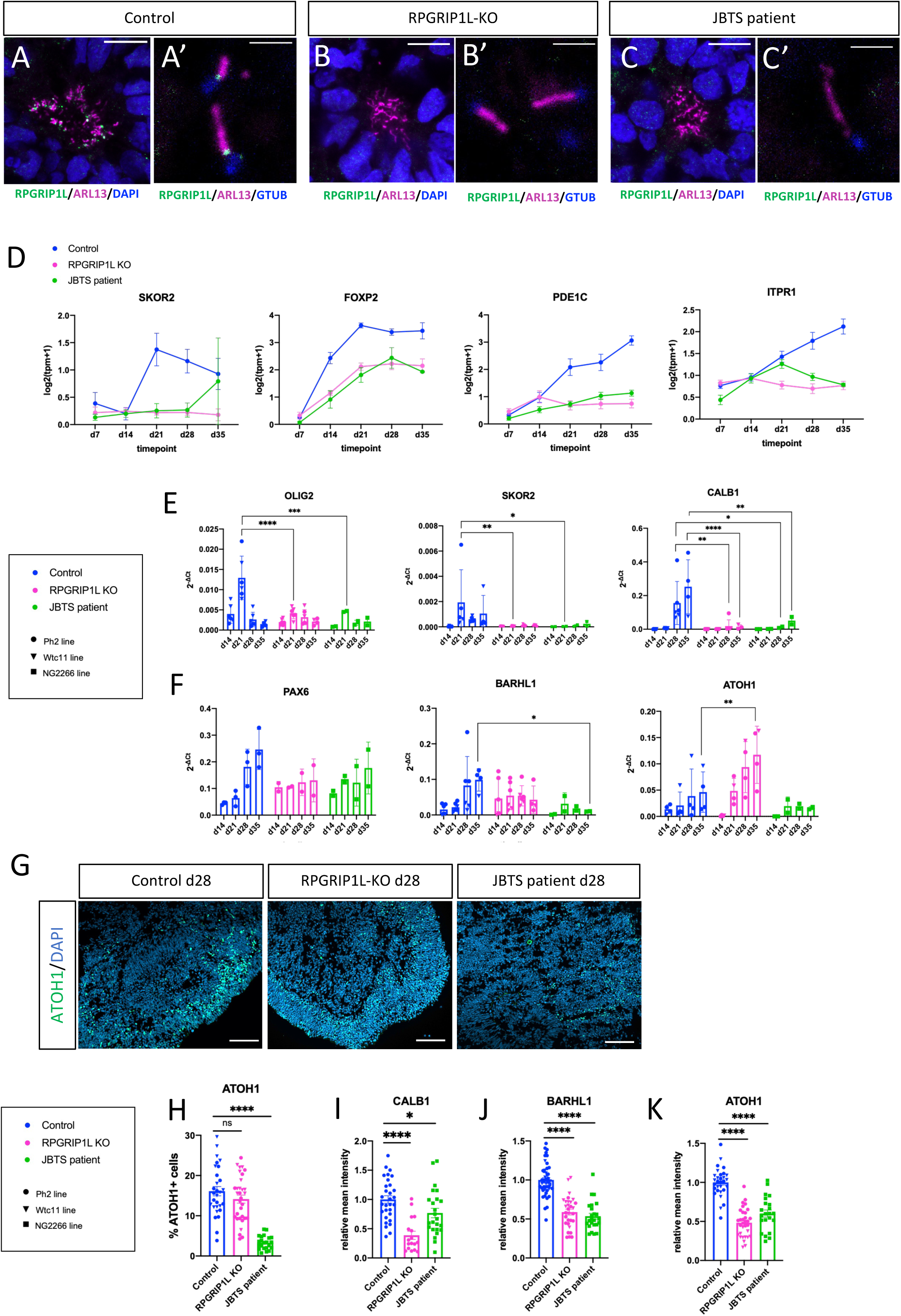
Altered RNA expression of Purkinje cell and glutamatergic lineage markers in *RPGRIP1L-*deficient cerebellar organoids **(A-C’)** Immunofluorescence on cryosections from control (A, A’), *RPGRIP1L* KO (B, B’) and JBTS patient (C, C’) organoids at day 21, labeling RPGRIP1L (in green) and the ciliary marker ARL13B (in magenta). Nuclei are stained with DAPI in A,B,C (blue) and basal bodies are stained with GTUB in A’, B’, C’ (blue). Scale bars: 10 µm in A-C and 2 µm in A’-C’. **(D)** Graph showing normalized temporal gene expression analysis from bulk RNAseq data, displayed as mean ± SEM, of the Purkinje lineage markers *SKOR2, FOXP2, PDE1C* and *ITPR1* along the differentiation protocol. For control and *RPGRIP1L* KO conditions data were obtained from both Ph2 and Wtc11 cell lines. See Supplementary Table 2 for number of independent experimental replicates. **(E-F)** qPCR analysis of (E) the PC markers *OLIG2, SKOR2* and *CALB1* and (F) the glutamatergic lineage markers *PAX6, BARHL1, ATOH1*, in cerebellar organoids at different timepoints from several independent experiments. Each dot in the graph represents one independent experiment per timepoint and condition; circular dots represent Ph2 line, triangular dots represent Wtc11 line, squared dots represent NG2266 line. Mean ± SEM are shown. Asterisks denote statistical significance according to ordinary Two-Way ANOVA (*P<0.033, **P < 0.0021, ***P<0.0002 and ****P< 0.0001). **(G)** Immunofluorescence for ATOH1 (glutamatergic lineage) on cryosections from control, *RPGRIP1L* KO and JBTS patient organoids at day 28. **(H)** Graph showing the % of ATOH1-positive cells over the total number of nuclei stained with DAPI. Positive cells were counted in more than 15 sections from different organoids per experiment per condition. Control and *RPGRIP1L* KO: N=3 (Ph2 N=2 – circular dots, Wtc11 N=1 – triangular dots); JBTS patient: NG2266 N=2 – squared dots. Each dot in the graph represent one organoid. Mean ± SEM are indicated. Asterisks denote statistical significance according to One-Way ANOVA Kruskal-Wallis test with Dunn’s correction (****P<0.0001). **(I-K)** Graphs showing the average intensity of CALB1 (I), BARHL1 (J), ATOH1 (K) labeling on cryosections from control, *RPGRIP1L* KO and JBTS organoids at day 28. Average cell intensity was measured in more than 15 sections from different organoids per experiment per condition. Relative mean intensity was calculated considering the mean intensity in control organoid sections as the reference value (1). Each dot in the graph represent one organoid. Mean ± SEM are indicated;. Asterisks denote statistical significance according to One-Way ANOVA Kruskal-Wallis test with Dunn’s correction (*P<0.033, and ****P<0.0001). N: number of independent experimental replicates. Scale bars: 100 µm in H.

**Supplementary Figure 3.**
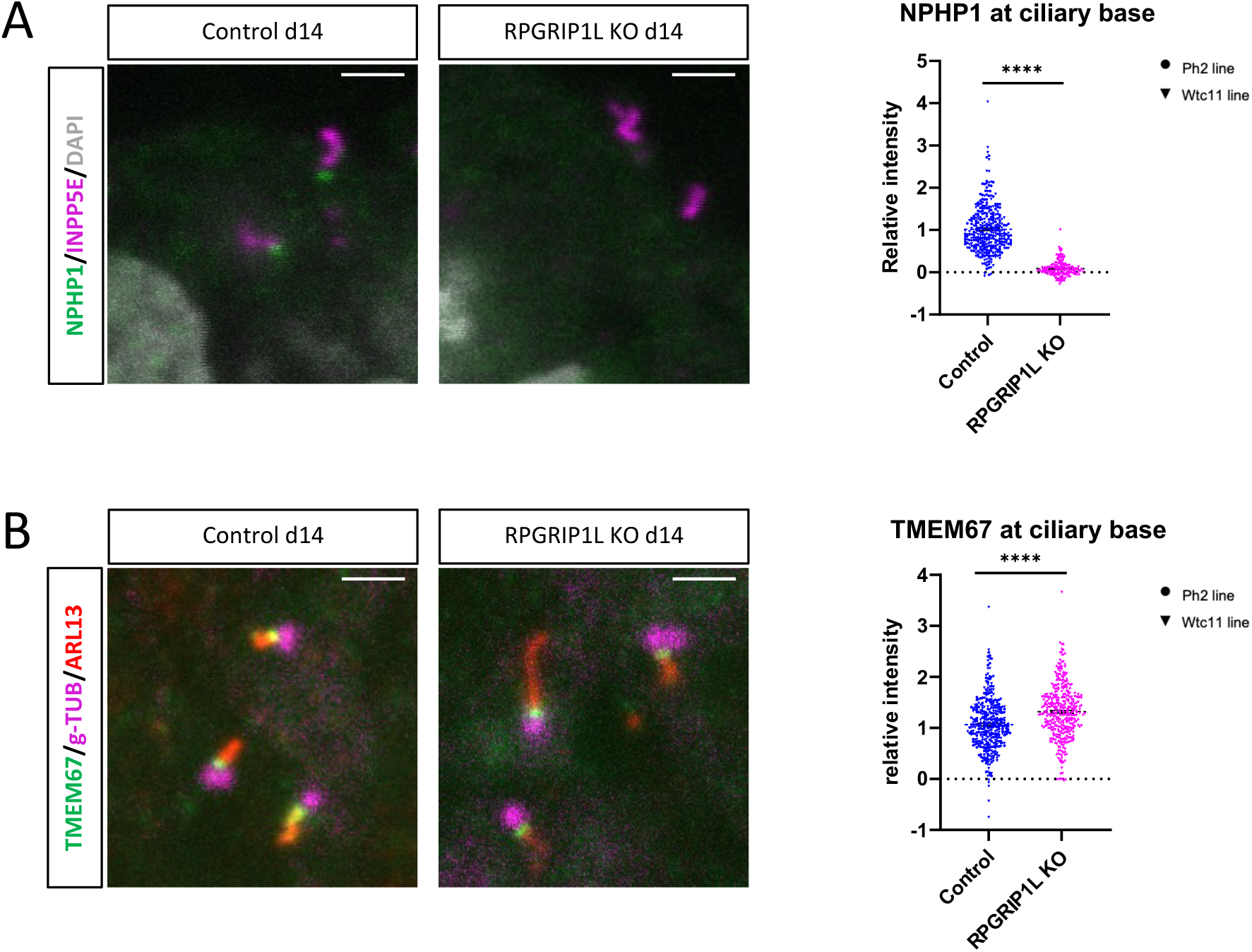
: Proteins from different TZ complexes respond differently to the loss of RPGRIP1L in day 14 cerebellar organoids. **(A)** Left: Immunofluorescence for NPHP1 (green) and INPP5E (magenta) on sections from control (left) and *RPGRIP1L* KO (right) organoids at day 14. DAPI is in white. Scale bar, 2 µm. Right: Graph showing the quantification of NPHP1 intensity at the TZ. Measures for individual cilia are represented and the mean value is indicated. Asterisks: statistical significance according to unpaired t test with Welch’s correction (****P < 0.0001)**. (B)** Left: Immunofluorescence for TMEM67 (green), ARL13B (red) and gamma Tubulin (g-TUB) (magenta) on sections from control (left) and *RPGRIP1L* KO (right) organoids at day 14. Scale bar, 2 µm. Right: Graph showing the quantification of TMEM67 intensity at the TZ. Measures for individual cilia are represented and the mean value is indicated. Asterisks: statistical significance according to unpaired t test with Welch’s correction (****P < 0.0001).

**Supplementary Figure 4.**
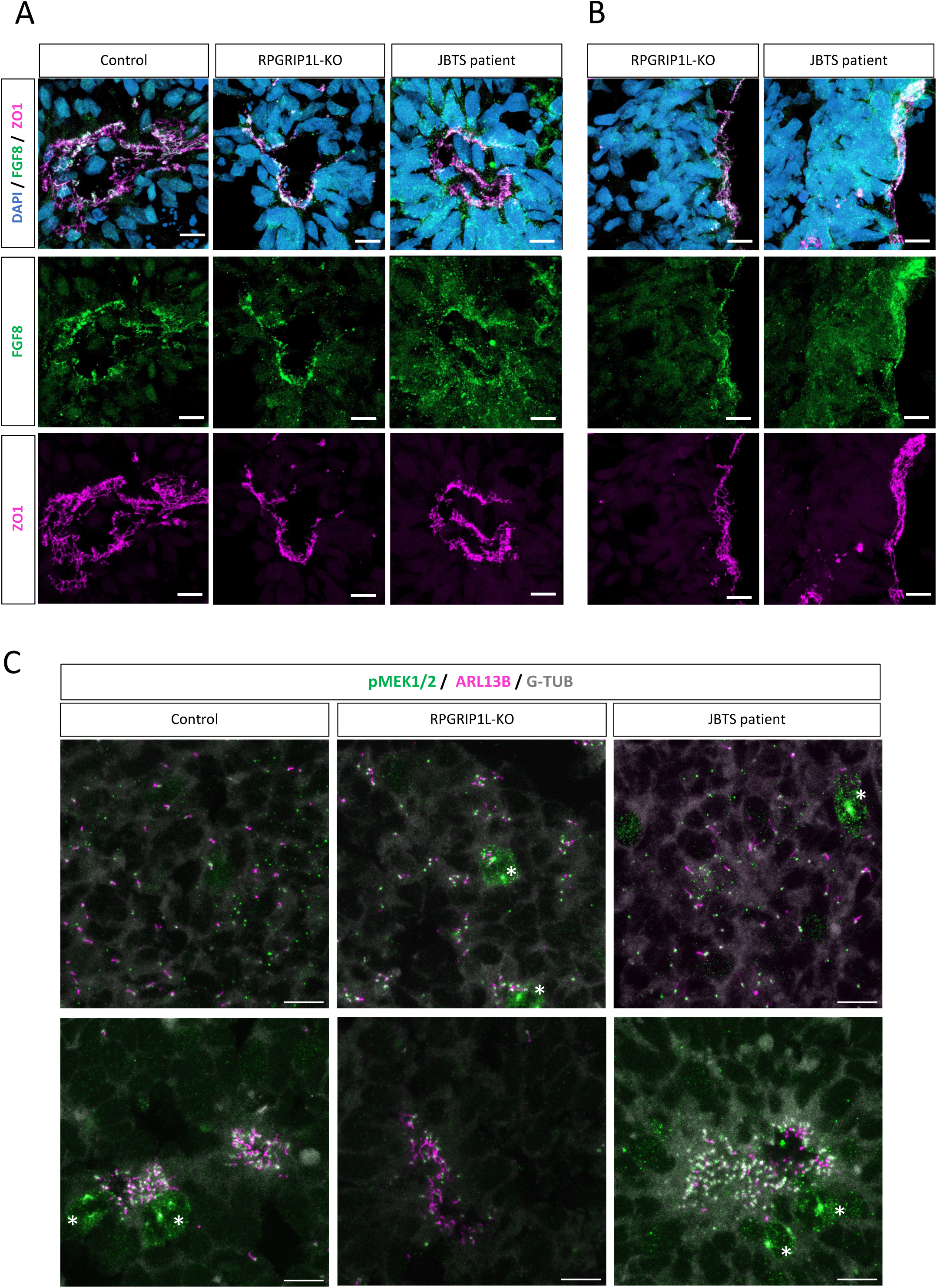
**FGF8 and P-Mek1/2 localization in early neural progenitors**

**Supplementary Figure 5.**
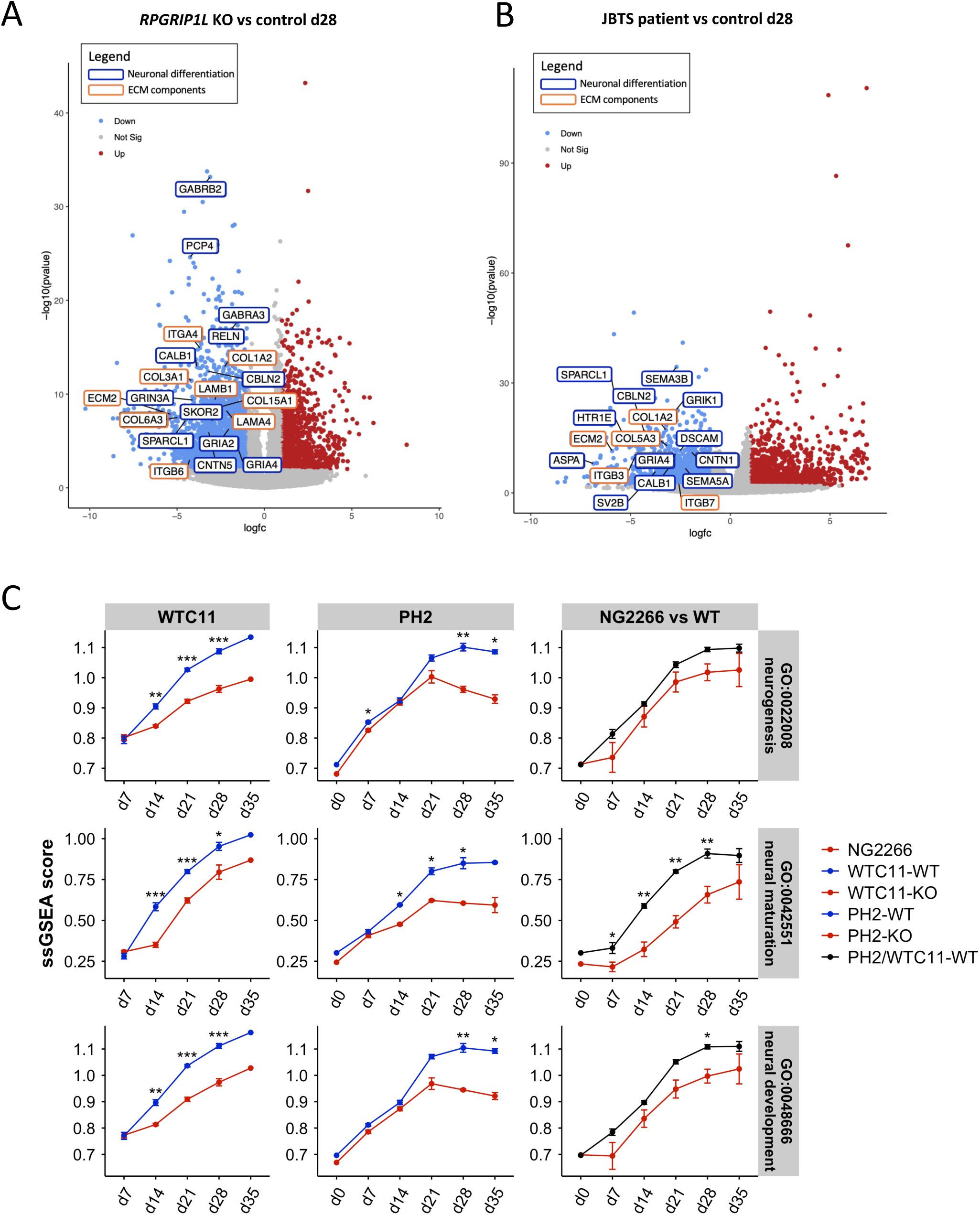
Further analysis of DEGs confirms altered neurogenesis and ECM production in RPGRIP1L-deficient organoids compared to controls **(A-B)** Volcano plots representing DEGs resulting from DESeq2 analysis (only genes with P<0.001, |log2FC|>2 are colored) in *RPGRIP1L* KO versus control condition (A) or in JBTS patient versus control condition (B) at day 28. Selected genes involved in neuronal differentiation or in building ECM are circled with different colors as indicated in the legend on the top right of the panel. **(C)** Graphs showing ssGSEA of neurogenesis and neuronal maturation GO terms in *RPGRIP1L* KO versus control organoids at different stages of differentiation.

**Supplementary Figure 6.**
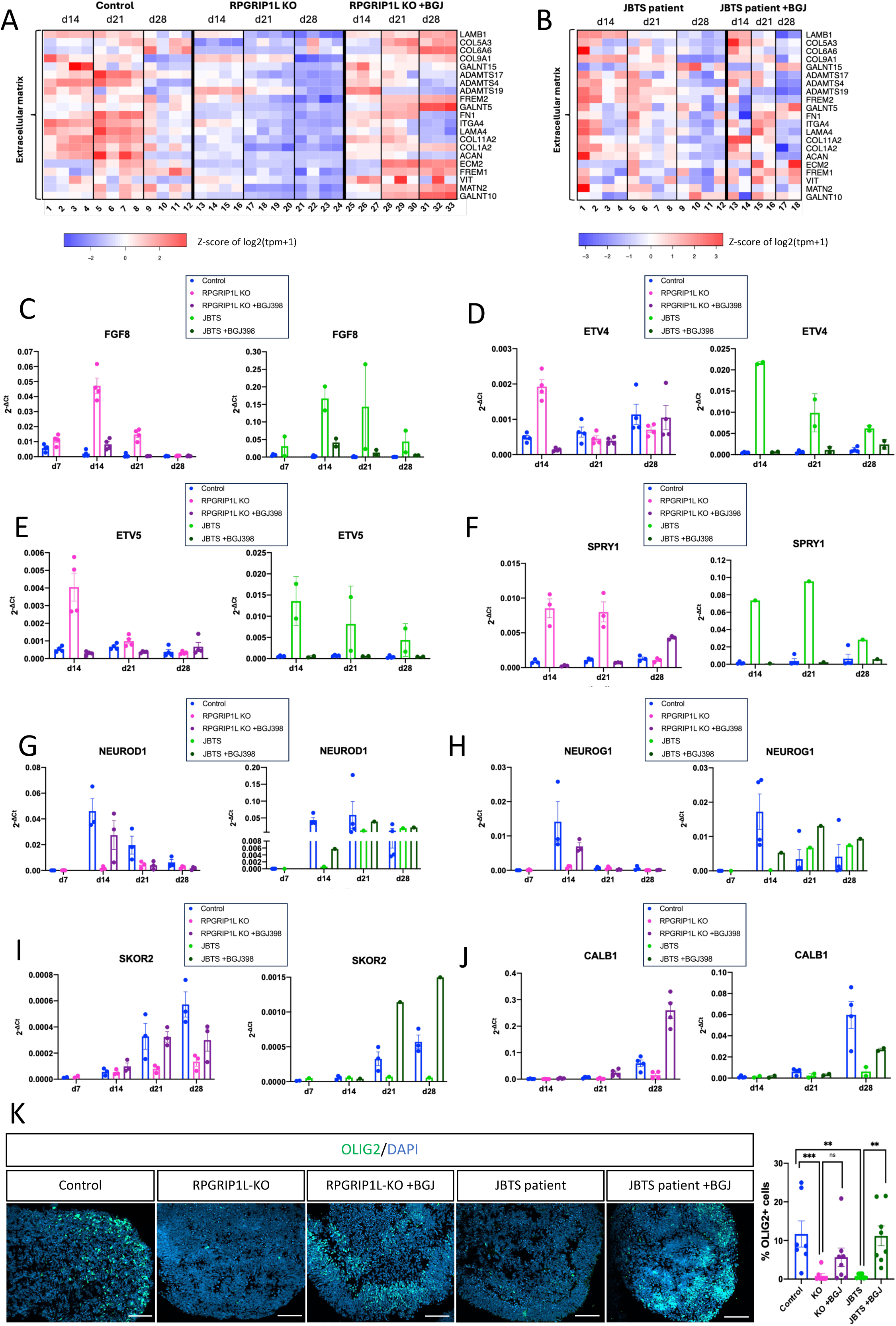
FGF signaling inhibition rescues organoid size and gene expression in *RPGRIP1L* deficient organoids. **(A-B)** Heatmap representation of the normalized gene expression values of selected differentially expressed genes in control and in *RPGRIP1L* KO cerebellar organoids untreated or treated with BGJ-398 (A) and JBTS patient-derived cerebellar organoids untreated or treated with BGJ-398 (B). See supplementary Table 3 for number of independent experiments and sample numbering in A and see supplementary Table 4 for number of independent experiments and sample numbering in B. **(C-J)** Graphs showing gene expression levels at different timepoints by qPCR analysis of *FGF8* (C), *ETV4* (D) *ETV5* (E)*, SPRY1* (F)*, NEUROD1* (G)*, NEUROG1* (H)*, SKOR2* (I) and *CALB1* (J) in control and *RPGRIP1L*-deficient cerebellar organoids untreated or treated with BGJ-398. Data are shown as individual values from independent experiments, mean ± SEM are indicated. For (C-E, J) control and *RPGRIP1L* KO: N=4 (Wtc11); JBTS patient: N=2 (NG2266). For (F-I) control and *RPGRIP1L* KO: N=3 (Wtc11); JBTS patient: N=1 (NG2266). N: number of independent experimental replicates. **(K)** Left: Immunofluorescence for OLIG2 (green) on cryosections from day 21 control and *RPGRIP1L*-deficient organoids untreated or treated with BGJ-398. Nuclei are stained with DAPI (blue). Scale bar, 100 µm. Right: Graph showing the % of OLIG2+ over DAPI+ nuclei, counted in several sections from different organoids per condition. Controls and *RPGRIP1L* KO: N=1 (Ph2 N=1); JBTS: NG2266 N=1. Each dot in the graph represents one organoid; mean ± SEM are shown. Asterisks denote statistical significance according to One-Way ANOVA Kruskal-Wallis test with Dunn’s correction (**P<0.0021; ***P<0.0002). N: number of independent experimental replicates.

**Supplementary Figure 7.**
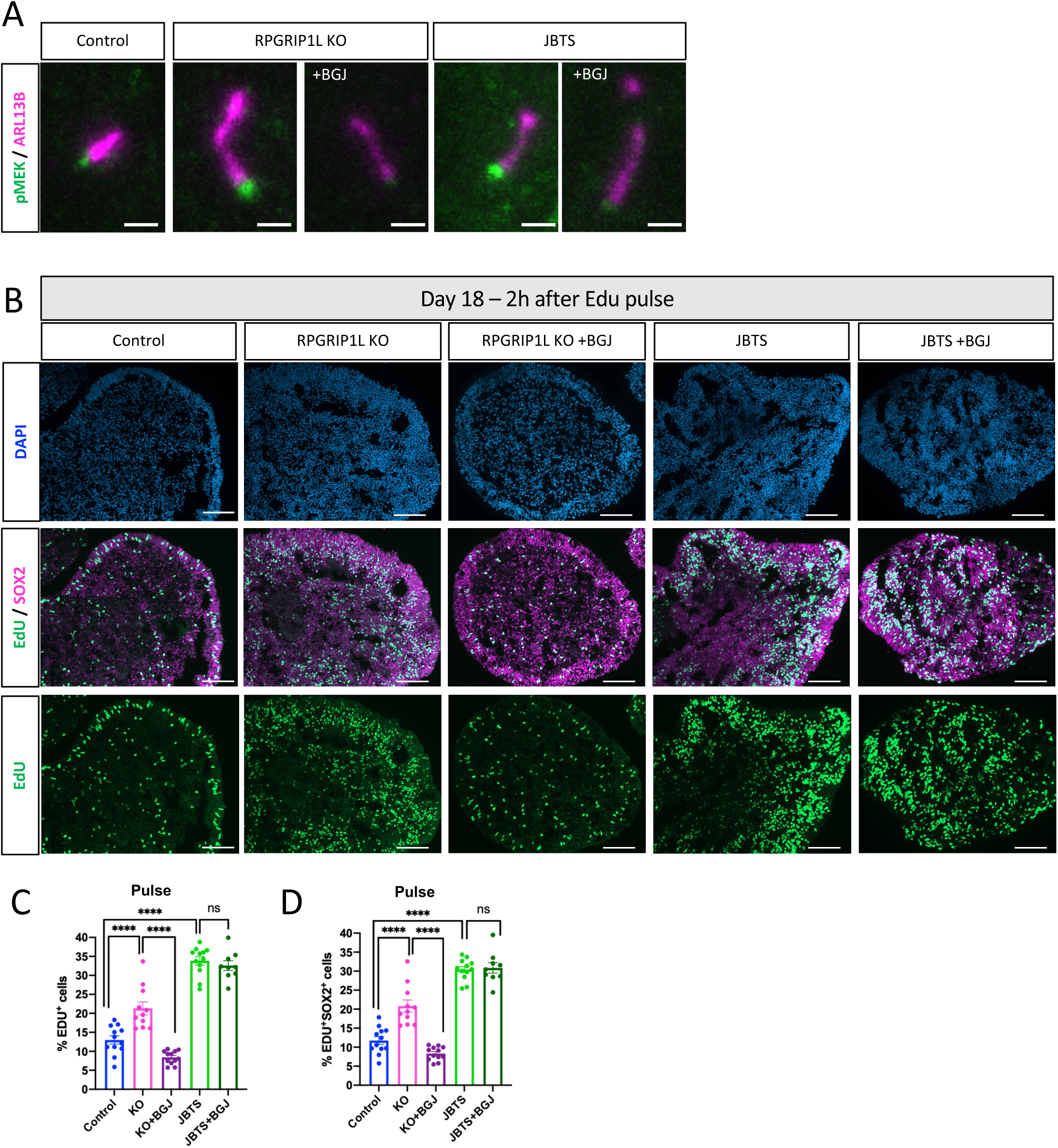
pMEK1/2 amount at the cilium base and neural progenitor proliferation are increased in *RPGRIP1L*-deficient organoids and partially rescued by blocking FGF signaling. **(A)** Immunofluorescence for pMEK1/2 (green) and ARL13B (magenta) (a marker of the ciliary axoneme) on organoid cryosections of the indicated control and RPGRIP1L-deficient genotypes, treated or not with BGJ-398. Scale bar, 1 µm **(B)** Immunofluorescence for SOX2 (magenta) and EdU (green) on organoid cryosections at day 18 – 2h after EdU pulse, from control and *RPGRIP1L*-deficient organoids, untreated or treated with BGJ-398. Scale bar, 100 µm **(C, D)** Graphs showing the % of EdU+ cells over the DAPI+ nuclei (C) and the % of EdU+SOX2+ cells over the total number of SOX2+ cells (D) at day 18 – 2h after EdU pulse, counted in several sections from different organoids per condition. Control and *RPGRIP1L* KO: N=1 (Wtc11); JBTS: N=1 (NG2266). Each dot in the graph represents one organoid; mean ± SEM are shown. Asterisks: statistical significance according to Ordinary-Way ANOVA after normal distribution testing (*P<0.033; ****P<0.0001).

**Supplementary Figure 8.**
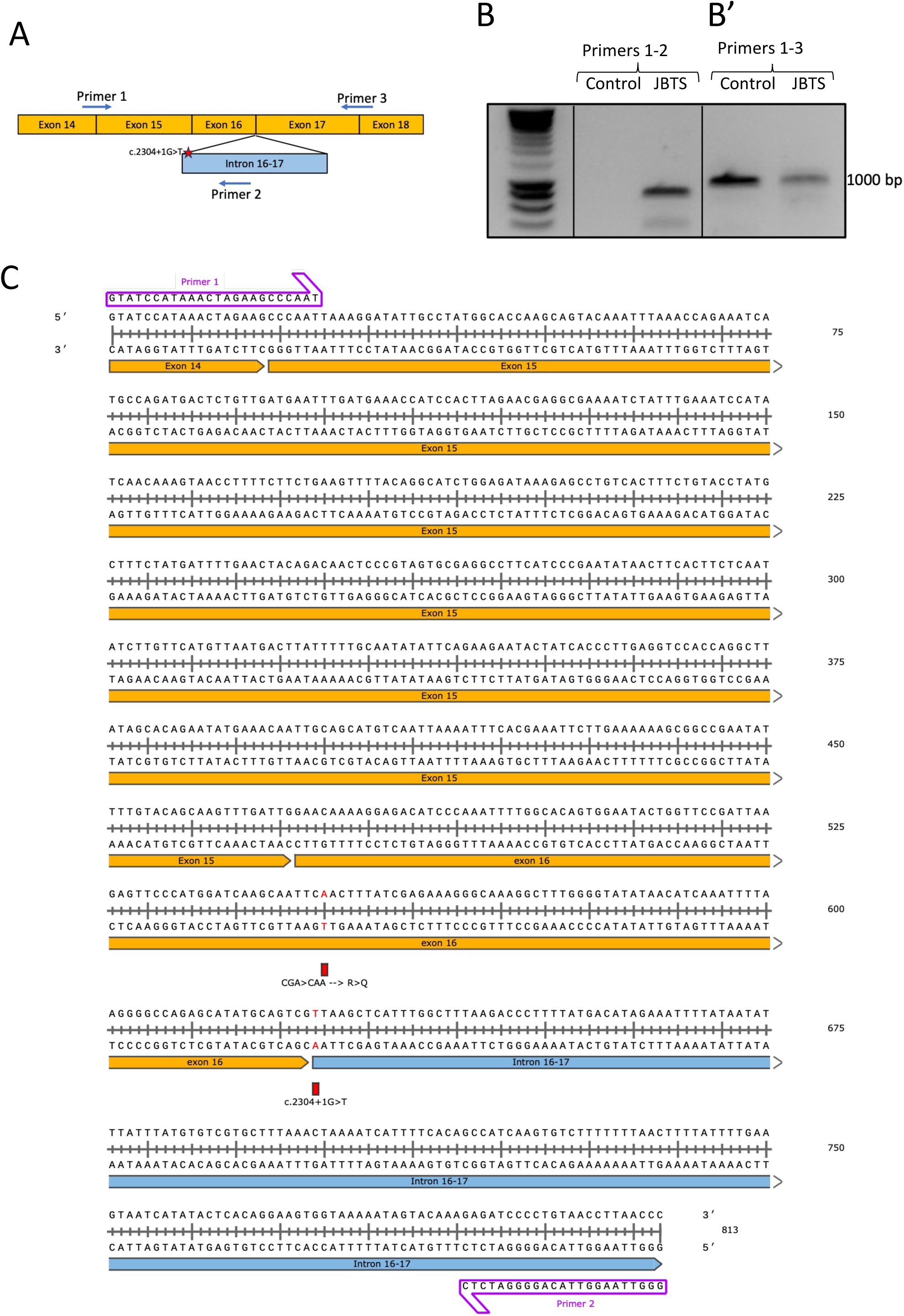
NG2266 splice variant c.2304+1G>T results in intron retention. **(A)** Schematic representation of *RPGRIP1L* gene exons 14-18 and PCR primers position to analyze cDNA products of *RPGRIP1L* gene in JBTS patient cells (NG2266 line) compared to control (Ph2 line). **(B)** PCR products from JBTS patient and control cerebellar cDNAs, templated by primer pairs as indicated, were analyzed by agarose gel electrophoresis and visualized with ethidium bromide. The primer pair 1-2 only amplifies JBTS patient cDNA, showing intron 16-17 retention. The primer pair 1-3 amplify both the second JBTS patient allele (encoding the *RPGRIP1L* Q684X variant) and control cDNA. **(C)** Sequence of the PCR product templated by primer pair 1-2 on cDNA derived from NG2266 cell line.

**Supplementary Table 1.** .Sample list Figure 1C

| **Label** | **Condition** | **N per timepont** | **Cell line** | **Sample name** |
|---|---|---|---|---|
| 1 | Control | Day 7 N=6 | Ph2 cell line | CBI-ph2-25-WT-d7 |
| 2 |  |  |  | CBII-ph2-25-WT-d7 |
| 3 |  |  | wtc11 cell line | CBIV-wtc11-19-WT-d7 |
| 4 |  |  |  | CBV-wtc11-19-WT-d7 |
| 5 |  |  |  | CBVI-wtc11-19-WT-d7 |
| 6 |  |  |  | CBVII-wtc11-19-WT-d7 |
| 7 | RPGRIP1L KO | Day 7 N=6 | Ph2 cell line | CBI-ph2-44-KO-d7 |
| 8 |  |  |  | CBII-ph2-44-KO-d7 |
| 9 |  |  | wtc11 cell line | CBIV-wtc11-65-KO-d7 |
| 10 |  |  |  | CBV-wtc11-65-KO-d7 |
| 11 |  |  |  | CBVI-wtc11-65-KO-d7 |
| 12 |  |  |  | CBVII-wtc11-65-KO-d7 |
| 13 | JBTS patient | Day 7 N=4 | NG2266 JBTS patient derived cell line | CBI-NG-2266-d7 |
| 14 |  |  |  | CBII-NG-2266-d7 |
| 15 |  |  |  | CBVII-NG-2266-d7 |
| 16 |  |  |  | CBVIII-NG-2266-d7 |

**Supplementary Table 2.** Sample list Figure 1E,F – 4D – 5A

| Label | Condition | N per timepont | Cell line | Sample name |
|---|---|---|---|---|
| 1 | Control | Day 7 N=6 | Ph2 cell line | CBI-ph2-25-WT-d7 |
| 2 |  |  |  | CBII-ph2-25-WT-d7 |
| 3 |  |  | wtc11 cell line | CBIV-wtc11-19-WT-d7 |
| 4 |  |  |  | CBV-wtc11-19-WT-d7 |
| 5 |  |  |  | CBVI-wtc11-19-WT-d7 |
| 6 |  |  |  | CBVII-wtc11-19-WT-d7 |
| 7 |  | Day 14 N=7 | Ph2 cell line | CBI-ph2-25-WT-d14 |
| 8 |  |  |  | CBII-ph2-25-WT-d14 |
| 9 |  |  |  | CBIII-ph2-25-WT-d14 |
| 10 |  |  | wtc11 cell line | CBIV-wtc11-19-WT-d14 |
| 11 |  |  |  | CBV-wtc11-19-WT-d14 |
| 12 |  |  |  | CBVI-wtc11-19-WT-d14 |
| 13 |  |  |  | CBVII-wtc11-19-WT-d14 |
| 14 |  | Day 21 N=7 | Ph2 cell line | CBI-ph2-25-WT-d21 |
| 15 |  |  |  | CBII-ph2-25-WT-d21 |
| 16 |  |  |  | CBIII-ph2-25-WT-d21 |
| 17 |  |  | wtc11 cell line | CBIV-wtc11-19-WT-d21 |
| 18 |  |  |  | CBV-wtc11-19-WT-d21 |
| 19 |  |  |  | CBVI-wtc11-19-WT-d21 |
| 20 |  |  |  | CBVII-wtc11-19-WT-d21 |
| 21 |  | Day 28 N=7 | Ph2 cell line | CBI-ph2-25-WT-d28 |
| 22 |  |  |  | CBII-ph2-25-WT-d28 |
| 23 |  |  |  | CBIII-ph2-25-WT-d28 |
| 24 |  |  | wtc11 cell line | CBIV-wtc11-19-WT-d28 |
| 25 |  |  |  | CBV-wtc11-19-WT-d28 |
| 26 |  |  |  | CBVI-wtc11-19-WT-d28 |
| 27 |  |  |  | CBVII-wtc11-19-WT-d28 |
| 28 |  | Day 35 N=4 | Ph2 cell line | CBI-ph2-25-WT-d35 |
| 29 |  |  |  | CBII-ph2-25-WT-d35 |
| 30 |  |  |  | CBIII-ph2-25-WT-d35 |
| 31 |  |  | wtc11 | CBIV-wtc11-19-WT-d35 |
| 32 | RPGRIP1L KO | Day 7 N=6 | Ph2 cell line | CBI-ph2-44-KO-d7 |
| 33 |  |  |  | CBII-ph2-44-KO-d7 |
| 34 |  |  | wtc11 cell line | CBIV-wtc11-65-KO-d7 |
| 35 |  |  |  | CBV-wtc11-65-KO-d7 |
| 36 |  |  |  | CBVI-wtc11-65-KO-d7 |
| 37 |  |  |  | CBVII-wtc11-65-KO-d7 |
| 38 |  | Day 14 N=6 | Ph2 cell line | CBI-ph2-44-KO-d14 |
| 39 |  |  |  | CBII-ph2-44-KO-d14 |
| 40 |  |  | wtc11 cell line | CBIV-wtc11-65-KO-d14 |
| 41 |  |  |  | CBV-wtc11-65-KO-d14 |
| 42 |  |  |  | CBVI-wtc11-65-KO-d14 |
| 43 |  |  |  | CBVII-wtc11-65-KO-d14 |
| 44 |  | Day 21 N=6 | Ph2 cell line | CBI-ph2-44-KO-d21 |
| 45 |  |  |  | CBII-ph2-44-KO-d21 |
| 46 |  |  | wtc11 cell line | CBIV-wtc11-65-KO-d21 |
| 47 |  |  |  | CBV-wtc11-65-KO-d21 |
| 48 |  |  |  | CBVI-wtc11-65-KO-d21 |
| 49 |  |  |  | CBVII-wtc11-65-KO-d21 |
| 50 |  | Day 28 N=6 | Ph2 cell line | CBI-ph2-44-KO-d28 |
| 51 |  |  |  | CBII-ph2-44-KO-d28 |
| 52 |  |  | wtc11 cell line | CBIV-wtc11-65-KO-d28 |
| 53 |  |  |  | CBV-wtc11-65-KO-d28 |
| 54 |  |  |  | CBVI-wtc11-65-KO-d28 |
| 55 |  |  |  | CBVII-wtc11-65-KO-d28 |
| 56 |  | Day 35 N=3 | Ph2 cell line | CBI-ph2-44-KO-d35 |
| 57 |  |  |  | CBII-ph2-44-KO-d35 |
| 58 |  |  | wtc11 | CBIV-wtc11-65-KO-d35 |
| 59 | JBTS patient | Day 7 N=4 | NG2266 JBTS patient derived cell line | CBI-NG-2266-d7 |
| 60 |  |  |  | CBII-NG-2266-d7 |
| 61 |  |  |  | CBVII-NG-2266-d7 |
| 62 |  |  |  | CBVIII-NG-2266-d7 |
| 63 |  | Day 14 N=4 |  | CBI-NG-2266-d14 |
| 64 |  |  |  | CBII-NG-2266-d14 |
| 65 |  |  |  | CBVII-NG-2266-d14 |
| 66 |  |  |  | CBVIII-NG-2266-d14 |
| 67 |  | Day 21 N=4 |  | CBI-NG-2266-d21 |
| 68 |  |  |  | CBII-NG-2266-d21 |
| 69 |  |  |  | CBVII-NG-2266-d21 |
| 70 |  |  |  | CBVIII-NG-2266-d21 |
| 71 |  | Day 28 N=4 |  | CBI-NG-2266-d28 |
| 72 |  |  |  | CBII-NG-2266-d28 |
| 73 |  |  |  | CBVII-NG-2266-d28 |
| 74 |  |  |  | CBVIII-NG-2266-d28 |
| 75 |  | Day 35 N=2 |  | CBI-NG-2266-d35 |
| 76 |  |  |  | CBII-NG-2266-d35 |

**Supplementary Table 3.** Sample list Figure 6C

| Label | Cell line | Condition | N per timepont | Sample name |
|---|---|---|---|---|
| 1 | wtc11 cell line | Control | Day 14 N=4 | CBIV-wtc11-19-WT-d14 |
| 2 |  |  |  | CBV-wtc11-19-WT-d14 |
| 3 |  |  |  | CBVI-wtc11-19-WT-d14 |
| 4 |  |  |  | CBVII-wtc11-19-WT-d14 |
| 5 |  |  | Day 21 N=4 | CBIV-wtc11-19-WT-d21 |
| 6 |  |  |  | CBV-wtc11-19-WT-d21 |
| 7 |  |  |  | CBVI-wtc11-19-WT-d21 |
| 8 |  |  |  | CBVII-wtc11-19-WT-d21 |
| 9 |  |  | Day 28 N=4 | CBIV-wtc11-19-WT-d28 |
| 10 |  |  |  | CBV-wtc11-19-WT-d28 |
| 11 |  |  |  | CBVI-wtc11-19-WT-d28 |
| 12 |  |  |  | CBVII-wtc11-19-WT-d28 |
| 13 |  | RPGRIP1L KO | Day 14 N=4 | CBIV-wtc11-65-KO-d14 |
| 14 |  |  |  | CBV-wtc11-65-KO-d14 |
| 15 |  |  |  | CBVI-wtc11-65-KO-d14 |
| 16 |  |  |  | CBVII-wtc11-65-KO-d14 |
| 17 |  |  | Day 21 N=4 | CBIV-wtc11-65-KO-d21 |
| 18 |  |  |  | CBV-wtc11-65-KO-d21 |
| 19 |  |  |  | CBVI-wtc11-65-KO-d21 |
| 20 |  |  |  | CBVII-wtc11-65-KO-d21 |
| 21 |  |  | Day 28 N=4 | CBIV-wtc11-65-KO-d28 |
| 22 |  |  |  | CBV-wtc11-65-KO-d28 |
| 23 |  |  |  | CBVI-wtc11-65-KO-d28 |
| 24 |  |  |  | CBVII-wtc11-65-KO-d28 |
| 25 |  | RPGRIP1L KO +BGJ398 treatment | Day 14 N=3 | CBV-wtc11-65-KO-BGJ-d14 |
| 26 |  |  |  | CBVI-wtc11-65-KO-BGJ-d14 |
| 27 |  |  |  | CBVII-wtc11-65-KO-BGJ-d14 |
| 28 |  |  | Day 21 N=3 | CBV-wtc11-65-KO-BGJ-d21 |
| 29 |  |  |  | CBVI-wtc11-65-KO-BGJ-d21 |
| 30 |  |  |  | CBVII-wtc11-65-KO-BGJ-d21 |
| 31 |  |  | Day 28 N=3 | CBV-wtc11-65-KO-BGJ-d28 |
| 32 |  |  |  | CBVI-wtc11-65-KO-BGJ-d28 |
| 33 |  |  |  | CBVII-wtc11-65-KO-BGJ-d28 |

**Supplementary Table 4.** Sample list Figure 6D

| Label | Cell line | Condition | N per timepont | Sample name |
|---|---|---|---|---|
| 1 | NG2266 JBTS patient derived cell line | NG2266 untreated | Day 14 N=4 | CBI-NG-2266-d14 |
| 2 |  |  |  | CBII-NG-2266-d14 |
| 3 |  |  |  | CBVII-NG-2266-d14 |
| 4 |  |  |  | CBVIII-NG-2266-d14 |
| 5 |  |  | Day 21 N=4 | CBI-NG-2266-d21 |
| 6 |  |  |  | CBII-NG-2266-d21 |
| 7 |  |  |  | CBVII-NG-2266-d21 |
| 8 |  |  |  | CBVIII-NG-2266-d21 |
| 9 |  |  | Day 28 N=4 | CBI-NG-2266-d28 |
| 10 |  |  |  | CBII-NG-2266-d28 |
| 11 |  |  |  | CBVII-NG-2266-d28 |
| 12 |  |  |  | CBVIII-NG-2266-d28 |
| 13 |  | NG2266 +BGJ398 treatment | Day 14 N=2 | CBVII-NG-2266-BGJ-d14 |
| 14 |  |  |  | CBVIII-NG-2266-BGJ-d14 |
| 15 |  |  | Day 21 N=2 | CBVII-NG-2266-BGJ-d21 |
| 16 |  |  |  | CBVIII-NG-2266-BGJ-d21 |
| 17 |  |  | Day 28 N=2 | CBVII-NG-2266-BGJ-d28 |
| 18 |  |  |  | CBVIII-NG-2266-BGJ-d28 |

**Supplementary Table 5.** Primary antibodies list

| **primary Antibody** | **host** | **company** | **reference** | **dilution** |
|---|---|---|---|---|
| Atoh1(Math1) | rabbit | Cohesion Biosciences | CQA1401 | 1:200 |
| Barhl1 | rabbit | Atlas | HPA004809 | 1:300 |
| Calbindin1 | mouse | Sigma | C9848 | 1:300 |
| Fgf8 | mouse IgG1 | ReD system | MAB323 | 1:40 |
| HuC/HuD (16A11) Elav | mouse IgG2b | Molecular Probes | A-21271 | 1:200 |
| Inpp5e | rabbit | Proteintech | 17797-1-AP | 1:300 |
| NeuroD1 | mouse IgG1 | Proteintech | 66691-1-Ig | 1:200 |
| Olig2 | rabbit | Millipore | AB9610 | 1:200 |
| pMEK1/2 | rabbit | Cell Signalling | 9121 | 1:100 |
| Sox2 | rabbit | Millipore | AB5603 | 1:200 |
| Sp8 | rabbit | Abcam | AB302916 | 1:200 |
| Zo1 | mouse IgG1 | Invitrogen | 33-9100 | 1:500 |
| *γ*-Tubulin | mouse IgG1 | Sigma | T6557 | 1:500 |

**Supplementary Table 6.** Secondary antibody list

| **secondary Antibody** | **host** | **conjugate** | **company** |
|---|---|---|---|
| Mouse IgG1 | goat | Alexa488 | Molecular Probes |
| Mouse IgG1 | goat | Alexa633 | Molecular Probes |
| Mouse IgG2b | goat | Alexa568 | Molecular Probes |
| Rabbit | goat | Alexa488 | Molecular Probes |
| Rabbit | goat | Alexa568 | Molecular Probes |
| Rabbit | goat | Alexa633 | Molecular Probes |

**Supplementary Table 7.**
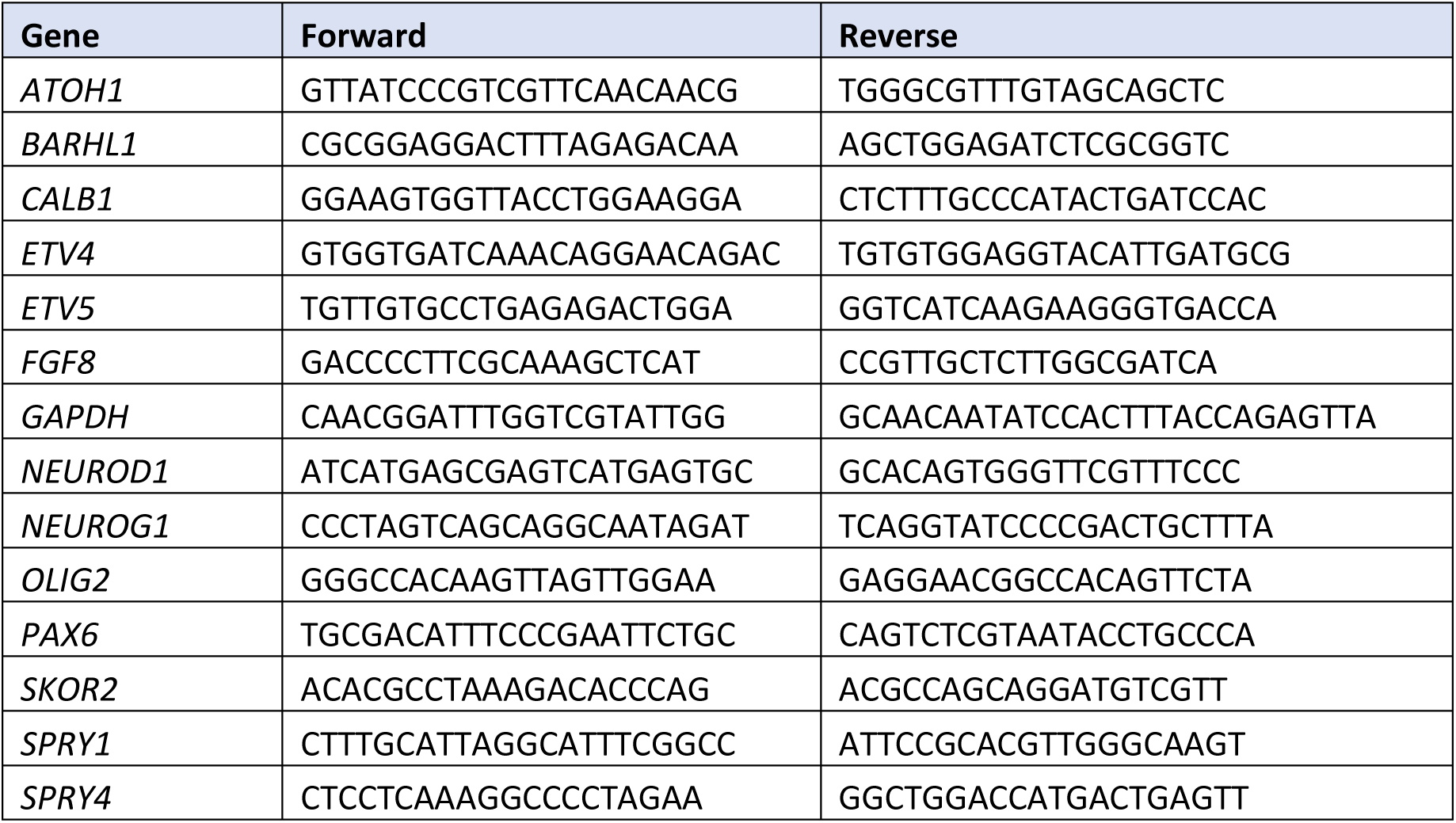
qPCR primer list

## Supplementary Excel Files

- Supplementary Excel file 1. Log2(tpm+1)_hiPSCs d0 and samples from Supplementary Tables 1-2
- Supplementary Excel file 2. DESeq2_table_KOvsWT_d14
- Supplementary Excel file 3. DESeq2_table_NG2266vsWT_d14
- Supplementary Excel file 4. DESeq2_table_KOvsWT_d28
- Supplementary Excel file 5. DESeq2_table_NG2266vsWT_d28
- Supplementary Excel file 6. GO_d28_downregulated-genes_KOvsWT
- Supplementary Excel file 7. GO_d28_downregulated-genes_JBTSvsWT
- Supplementary Excel file 8. Log2(tpm+1)_samples from Supplementary Table 3-4

